# Viscophobic turning dictates microalgae transport in viscosity gradients

**DOI:** 10.1101/2020.11.05.369801

**Authors:** Michael R. Stehnach, Nicolas Waisbord, Derek M. Walkama, Jeffrey S. Guasto

## Abstract

Gradients in fluid viscosity characterize microbiomes ranging from mucus layers on marine organisms^1^ and human viscera^2,3^ to biofilms^4^. While such environments are widely recognized for their protective effects against pathogens and their ability to influence cell motility^2,5^, the physical mechanisms regulating cell transport in viscosity gradients remain elusive^6–8^, primarily due to a lack of quantitative observations. Through microfluidic experiments, we directly observe the transport of model biflagellated microalgae (*Chlamydomonas reinhardtii*) in controlled viscosity gradients. We show that despite their locally reduced swimming speed, the expected cell accumulation in the viscous region^9,10^ is stifled by a viscophobic turning motility. This deterministic cell rotation – consistent with a flagellar thrust imbalance^11,12^ – reorients the swimmers down the gradient, causing their accumulation in the low viscosity zones for sufficiently strong gradients. Corroborated by Langevin simulations and a three-point force model of cell propulsion, our results illustrate how the competition between viscophobic turning and viscous slowdown ultimately dictates the fate of population scale microbial transport in viscosity gradients.

---

Microbial transport is driven by external cues^5,14^ and plays a crucial role in regulating the function of ecosystems ranging from cell colonies^4^ to Earth’s biome^15^. Swimming microorganisms navigate toward regions of optimal nutrient uptake and photosynthetic productivity by sensing gradients in chemical concentration (i.e. chemotaxis^3,16^) and light intensity (i.e. phototaxis^17^), respectively. External forces or torques acting upon cells can also mechanically guide microbes toward favorable conditions: Gravitational fields orient eukaryotic marine microbes within the water column through gravitaxis^18,19^, and magnetic fields serve as a compass for bacteria in sedimentary environments via magnetotaxis^20^. Additionally, swimming cells are widely known to respond to gradients in fluid viscosity^5^, which can vary broadly in scale and strength^21,22^. Mucus layers (≈1 mm thick^23,24^) exhibit viscosities as low as 2 cP in corals and sessile marine organisms^25^, while in mammals they can range from 5 cP in the trachea to >1,000 cP in the gastrointestinal tract^22^. Viscous mucus serves as the first line of defense against motile pathogenic bacteria in humans^2,26^ and other organisms^1^, and it sustains endangered corals by trapping and distributing microorganisms and nutrient particles^1^. Despite the far reaching importance of these systems, a definitive paradigm for swimming cell transport in viscosity gradients is yet to be established, and the physical mechanisms controlling viscotactic motility remain unresolved.

Contrary to the widely studied effects of elevated homogeneous viscosity on cell motility^13,27^, experimental observations of swimming cell transport in inhomogeneous viscous environments are severely lacking and largely inconclusive. In viscosity gradients of unspecified strength, qualitative observations of cell density showed positive^6,7^ and negative^8^ (i.e. viscophobic) viscotaxis toward high and low viscosity regions, respectively. Recent theoretical and numerical work on swimmer hydrodynamics^11,12,28^ suggests that tuning the cell propulsive apparatus or body asymmetry could produce swimming either up or down the gradient. While these early experiments and deterministic simulations have provided some insights, the stochastic nature of cell motility and accumulation mechanisms rooted in local swimming speed modulation were largely neglected^9,10^. A holistic understanding of cell transport in viscosity gradients and the potential mechanisms underlying viscotactic motility have been hampered by a dearth of direct quantitative experiments.

In this Letter, we provide direct evidence of viscophobic motility for the model, biflagellated, microalga *Chlamydomonas reinhardtii*^29,30^ (Fig. 1a), which is achieved through a comprehensive, experimental description of its transport in a prescribed, microfluidic viscosity gradient (Fig. 1b). Our experiments show that as the magnitude of the gradient is increased, a muted cell accumulation in the high viscosity region^9^ (Fig. 1c,d) gives way to strong accumulation in the low viscosity region (Fig. 1e). Analysis of cell trajectories and complementary Langevin simulations reveal that this surprising transport behavior results from the competition between viscous accumulation due to reduced swimming speed^9,10^ and a viscophobic reorientation mechanism. The latter is consistent with a physical, hydrodynamic phenomenon^12^, which results in an ensemble drift of swimming *C. reinhardtii* down the viscosity gradient. Viscophobic reorientation dynamics bear striking similarities to a range of other taxes^19,20^ and add to the rich diversity of microbial transport mechanisms that regulate important macroscopic ecological processes^1–4,14^.

**Fig. 1:**
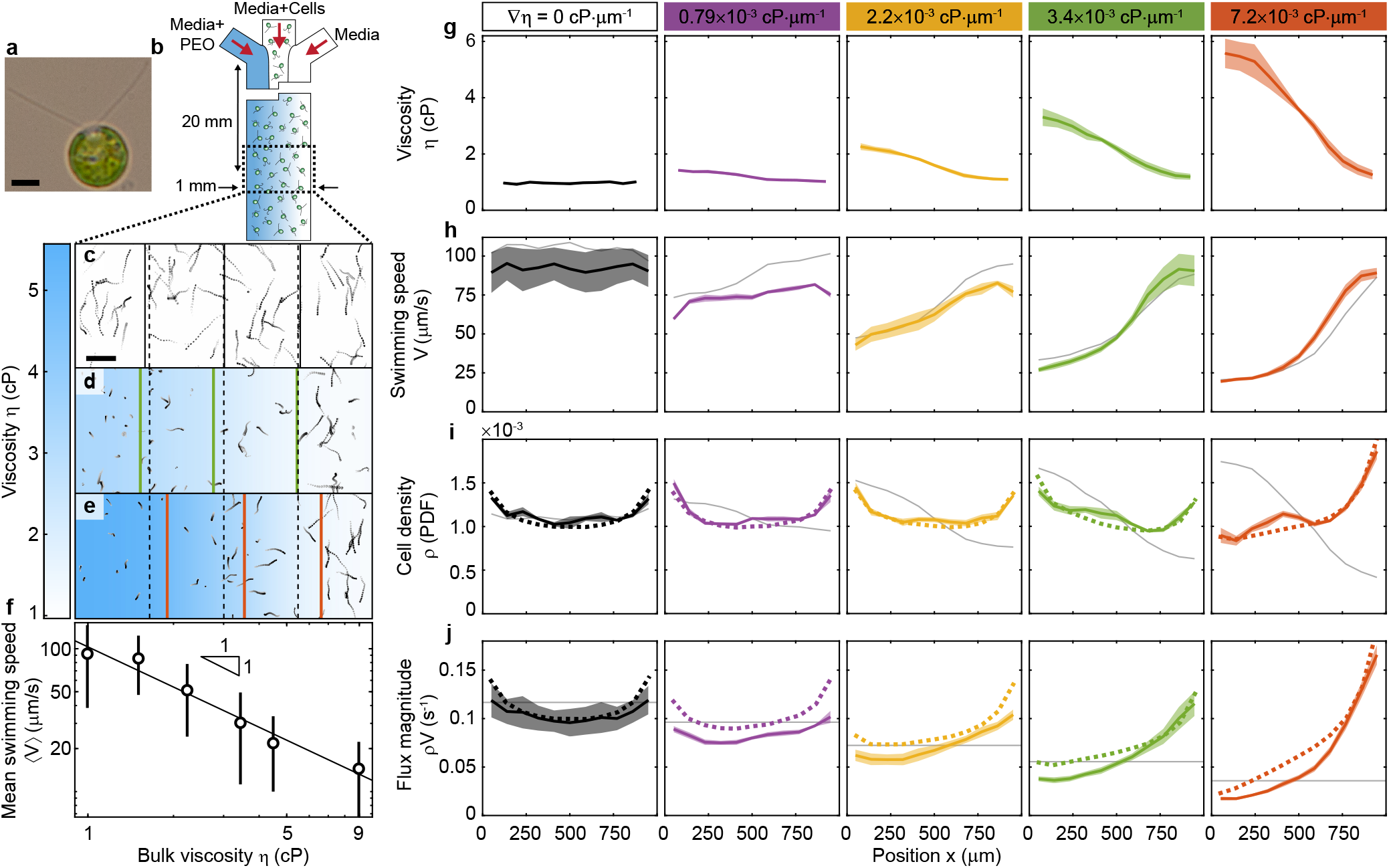
Viscosity gradient strength regulates dispersal of swimming microalgae. **a**, Biflagellated swimming microalga, *Chlamydomonas reinhardtii* (wild-type). Scale bar, 5 *μ*m. **b**, Schematic of three-inlet microfluidic device for viscosity gradient generation (100 *μ*m deep). Viscous media (PEO in M1 media), *C. reinhardtii* suspension, and M1 media are initially flow-stratified (Methods). Viscosity gradient is formed by polymer diffusion upon stopping flow (Extended Data Fig. 1). **c-e**, Multi-exposure images (1 s duration; intensity increases with time; 60 min post flow) of *C. reinhardtii* show a homogeneous distribution in uniform viscosity (**c**; ▽*η* = 0 cP·*μ*m^−1^, control), slight accumulation in high viscosity zone for moderate viscosity gradient (**d**; ▽*η* = 3.4 × 10^−3^ cP·*μ*m^−1^), and accumulation in low viscosity zone for large gradient (**e**; ▽*η* = 7.2 × 10^−3^ cP·*μ*m^−1^). Dashed black lines are 25%, 50%, and 75% of the channel width, and solid lines are the quartiles of cell density in **i**. Colorbar is measured viscosity profile from g. Scale bar, 100 *μ*m. **f**, Ensemble-averaged cell swimming speed, 〈*V*〉, in bulk viscous media, *η* (Supplementary Section 3.1; ≈2,700 trajectories each). Error bars are mean standard deviation (*N* = 2-3) and black line is a power-law fit^13^ (∝ *η*^−0.93^). **g**, Microfluidic viscosity profiles measured by microrheology (see Methods), *η*(*x*), characterized by the viscosity gradient magnitude, ▽*η* (Extended Data Figs. 1 and 8). Shaded area is standard deviation (30-90 min post flow). **h-j**, Measured profiles (solid colored curves) of local mean cell swimming speed (**h**), cell density (**i**), and flux magnitude (**j**), corresponding to **g**. Shaded regions indicate standard error (*N* = 3-4 with ≈2,100-9,200 trajectories each). Dashed curves are Langevin simulations with viscophobic turning (Fig. 3). Gray curves are analytical prediction for random walk swimmers^9^ without viscophobic turning (Supplementary Section 4.1).

To explain the observed migratory behavior of *C. reinhardtii*, microfluidic devices were designed to precisely control the viscosity gradient over length scales relevant to the swimming cells (Fig. 1b). Briefly, three solutions, including a Newtonian viscous polymer solution (polyethylene oxide, PEO), a dilute suspension of *C. reinhardtii*, and cell media, are stratified by continuous injection into the microchannel test section (Fig. 1b and Methods). Upon stopping the flow, the polymer diffuses across the strata for 30 min to form a smooth viscosity profile (Extended Data Fig. 1); thus providing ample time for swimming cells to disperse across the channel. Observations are performed between 30 and 90 min, during which time the viscosity profile, *η*(*x*), quantified by microrheology, is quasi-steady (Fig. 1g and Methods). In the absence of a viscosity gradient, their random walk motility yields a uniform cell density, *ρ*, across the channel (▽*η* = 0 cP·*μ*m^−1^; Fig. 1i), apart from minor accumulation at the boundaries due to the nature of cell-surface interactions^31^ (Supplementary Section 5.1.2).

For a directionally unbiased random walk, cell density is well known to scale inversely with local swimming speed^9,10,32^, *ρ*(*x*) ∝ 1/*V*(*x*). Imposing a moderate viscosity gradient (▽*η* = 3.4 × 10^−3^ cP·*μ*m^−1^) results in a slight accumulation of cells (Fig. 1i, solid colored lines) in the high viscosity region that is incommensurate with the strong local reduction in swimming speed (Fig. 1h, colored lines). The swimming speed distribution quantitatively agrees with analytical predictions (Fig. 1h, gray lines) based on the measured swimming speed of *C. reinhardtii* in homogeneous conditions^13^ (Fig. 1f) and the independently measured viscosity profile (Fig. 1g and Supplementary Section 4.1). However, this analytical prediction for the slowdown accumulation largely over-estimates^9,10^ the observed weakly positive viscotactic behavior (Fig. 1i, gray line). Stronger gradients reveal a counterintuitive viscophobic regime, where cells accumulate in the low viscosity region (Fig. 1i, ▽*η* = 7.2 × 10^−3^ cP·*μ*m^−1^). Moreover, the local cell flux magnitude, *ρ*(*x*)*V*(*x*), is expected to be spatially uniform across the channel for an unbiased random walk (Fig. 1j, gray lines). As ▽*η* increases, the progressive deviation of the measured flux magnitude (Fig. 1j, colored solid lines; Supplementary Section 4.1) from the predicted uniform value is a clear indication of an underlying viscophobic motility bias. To unravel these population scale observations and determine the nature of the observed behavior, we examine the kinematics of single cell motility in the viscosity gradients.

Cells swimming in a viscosity gradient exhibit directional dependence on their rotational kinematics and ultimately reorient down the viscosity gradient. In the absence of a viscosity gradient, cells generally maintain their initial swimming direction for several seconds (Fig. 2a). Conversely, cell trajectories initially oriented perpendicular (*θ* = ±*π*/2) to the gradient continuously bend toward the low viscosity region (*θ* = 0; Fig. 2b). This effect is made more apparent by examining the mean cell trajectory (Fig. 2b, thick colored curve): Due to the stochastic nature of microbial motility, for example stemming from noise in flagellar actuation and thermal fluctuations^33,34^ (persistence time, *τ*_p_ = 1/2*D*_r_ = 7.2 s; rotational diffusion coefficient, *D_r_*; see Methods), a large number of observations are necessary to unveil the underlying deterministic dynamics. Considering all swimming directions, the cells’ instantaneous angular velocity displays a sinusoidal dependence on orientation, which is absent from control experiments (Fig. 2c). We hypothesize that viscophobic turning in *C. reinhardtii* results from a slight viscosity difference experienced by its two flagella, causing an imbalance of their drag-based thrust forces^35^ (Fig. 2d, inset).

**Fig. 2:**
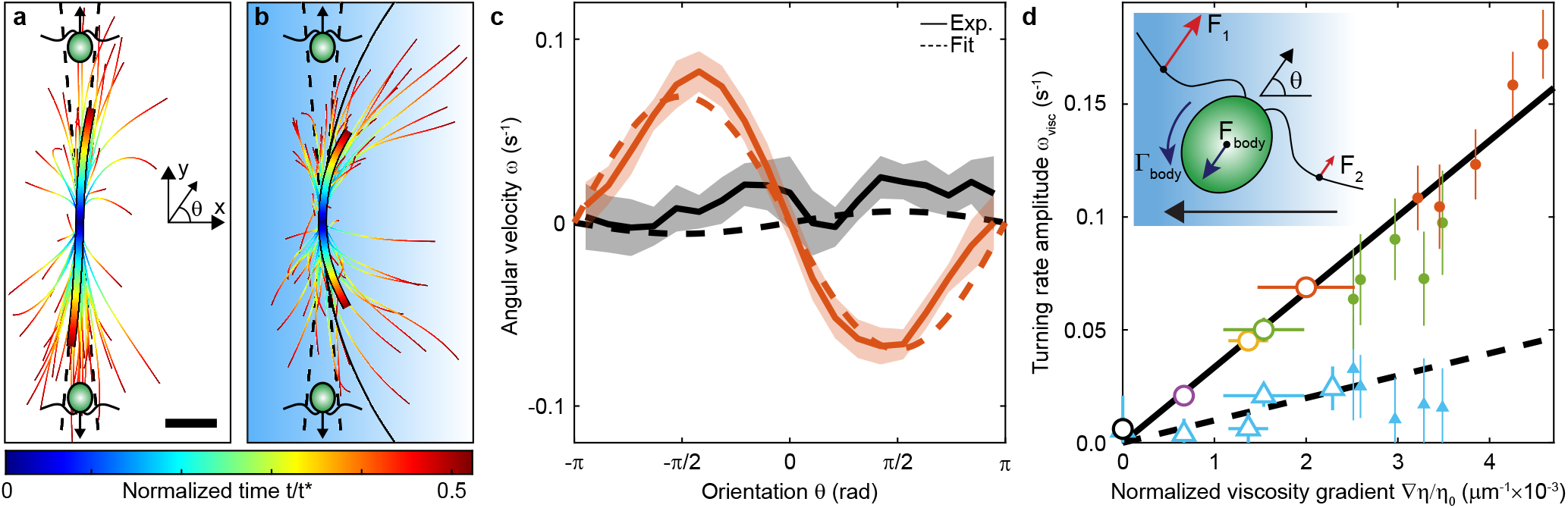
Coupling of swimming kinematics to viscosity gradient induces viscophobic turning. **a,b**, Measured wild-type cell trajectories for control (**a**; ▽*η* = 0 cP·*μ*m^−1^; 65 trajectories) and maximum viscosity gradient (**b**; ▽*η* = 7.2 × 10^−3^ cP·*μ*m^−1^; 76 trajectories). Conditionally sampled from 400 ≤ *x* ≤ 600 *μ*m and perpendicular to viscosity gradient (|*θ*| = π/2 ± 5°, dashed lines). Thick colored curve is mean trajectories, and black curve in **b** is analytical prediction (Supplementary Section 4.3). Background qualitatively indicates ▽*η*, and *t\** = 500 *μ*m/〈V(x)〉 is characteristic swimming time in channel. Scale bar, 100 *μ*m. **c**, Angular cell velocity exhibits orientation dependence in a viscosity gradient (▽*η* = 7.2 × 10^−3^ cP·*μ*m^−1^, orange) compared to control (▽*η* = 0 cP·*μ*m^−1^, black). Dashed lines are sinusoidal fits, and shaded region is standard error (7,730 and 3,454 trajectories for pooled replicates of black and orange, respectively). **d**, Viscophobic turning rate amplitude, *ω*_visc_, (**c**) increases with normalized viscosity gradient, ▽*η*/*η*_0_. Wild-type cells (open circles; 3,454 to 7,730 trajectories each) exhibit stronger viscophobic turning compared to short-flagella mutants (open triangles; 792-3,859 trajectories each) in quasi-steady conditions (30-90 min post flow). Solid and dashed black lines are linear fits to open circles and open triangles, respectively. Filled markers correspond to strong gradients sampled from transient viscosity profiles at early times (5-25 min post flow; 319-760 trajectories each; see also Extended Data Fig. 1 and 9). Filled blue triangles are transient times corresponding to ▽*η* = 3.4 × 10^−3^ cP·*μ*m^−1^ quasi-steady experiments. Horizontal error bars stem from measured viscosity profile, and vertical error bars (obscured by markers) are from fit in c. Inset: Potential mechanism for orientation-dependent viscophobic turning via propulsive force imbalance between flagella (Supplementary Section 4.2). Background and arrow indicate increasing viscosity gradient.

A three-point force swimmer model (Supplementary Section 4.2) quantitatively predicts the sinusoidal form of the observed cell rotation rate (Fig. 2c, dashed curves) and captures the mean swimming trajectory of the cells (Fig. 2b, black curve). The angular cell velocity is well fit by the model prediction (Fig. 2c, dashed curves), *ω*(*θ*) = −*ω*_visc_ sin(*θ*), and quantifies the turning rate amplitude, *ω*_visc_. The amplitude grows continuously with the normalized viscosity gradient, ▽*η*/*η*_0_, where *η*_0_ is the viscosity in the channel center, and the largest measured quasi-steady gradient yields a turning time, *τ* = 1/*ω*_visc_ ≈ 14 s (Fig. 2d). Furthermore, the linear dependence^12^ on ▽*η*/*η*_0_ (Fig. 2d, black line) stems from the growth of the turning rate with the gradient ▽*η*, which is tempered by enhanced rotational drag through increasing *η*_0_. A fit of the model prediction (Fig. 2d, black line; Supplementary Section 5.2), *ω*_visc_ = (*V*\*/2)▽*η*/*η*_0_, gives *V*\* ≈ 66 *μ*m/s, which is comparable to the swimming speed of *C. reinhardtii* in the tested gradients. The cell turning dynamics were also examined during early times in the viscotaxis assay (5-25 min post flow), when the viscosity gradient is rapidly evolving (Fig. 2d, filled circles). This analysis revealed that the linear increase of *ω*_visc_ persists at much higher gradients and emphasizes that ▽*η*/*η*_0_ controls the growth. Viscophobic reorientation of *C. reinhardtii* appears to be deterministic and omnipresent in viscosity gradients, but the magnitude and nature of the response is also expected to be a strong function of swimmer morphology. For biflagellated *C. reinhardtii*, the viscophobic turning rate amplitude decreases with flagellar length. A viscophobic aligning torque is generated by an imbalance in flagellar thrust, whose location, *d*, is a fraction of the flagellar length, *ℓ* To illustrate the dependence on flagellar length, *ω*_visc_ was measured for *C. reinhardtii* mutants, whose flagella are 36% shorter compared to wild-type cells^36^. The short-flagella cells (Fig. 2d, triangles) exhibit an approximately linear increase in *ω*_visc_ that is significantly diminished by a factor 3.3 compared to wild-type cells. Beyond other recent theoretical developments^11,12^ (Fig. 2d, dashed black line), our three-point force model uniquely predicts that flagellar length also determines whether the viscotactic response is negative or positive. The viscophobic aligning torque due to the flagellar thrust imbalance competes with an opposing torque due to a drag imbalance on the spheroidal cell body (i.e., translational-rotational coupling^37^). Biflagellates are thus predicted to behave in a viscophobic manner, when the lateral flagellar center of thrust is sufficiently large 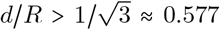, compared to the cell body radius, *R*. Conservatively, taking^29^ *R* ≈ 5 *μ*m and *d* ≈ *ℓ*/2, both wild-type cells (*d*/*R* ≈ 1.1) and short-flagella mutants (*d*/*R* ≈ 0.7) are expected to exhibit negative viscostaxis, exemplified by our experiments (Fig. 2d). The strong agreement of our measurements with our predictive model and the consistency with other recent models^11,12^ points toward a hydrodynamic origin for viscotaxis.

While viscophobic turning appears rooted in the complexity of swimmer hydrodynamics, it results in strikingly simple reorientation dynamics, which comprise a broader, universal class of motilities. The time evolution of the measured cell orientation distribution, p(*θ*|*t*), illustrates the condensation of the cell swimming direction down the viscosity gradient (Fig. 3a, top and Extended Data Fig. 2). Similar to most microbes, the random walk nature of *C. reinhardtii* motility stems from noise in their orientation^33^, modeled as rotational diffusion superimposed on the deterministic viscophobic turning:

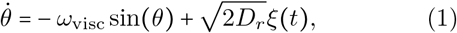

where *ξ* is Gaussian noise. A Langevin simulation (see Methods) accurately reproduces the time evolution of cell orientation (Fig. 3a, bottom and Extended Data Fig. 2), which forms a well-defined peak after several seconds (Fig. 3b). Our observations of viscotactic turning are thus consistent with deterministic responses that underlie most eukaryotic systems^38^, as opposed to the stochastic navigation of bacteria^16^. Importantly, viscotaxis appears to share universal dynamics that underpin a host of microbial responses to their physical environments – including gravitaxis^18,19^ and magnetotaxis^20,39^ – which are governed by a sinusoidal, paramagnetic-like torque in response to an external field. However, viscophobic reorientation is distinctive in that self-propulsion is fundamental to the cell response.

**Fig. 3:**
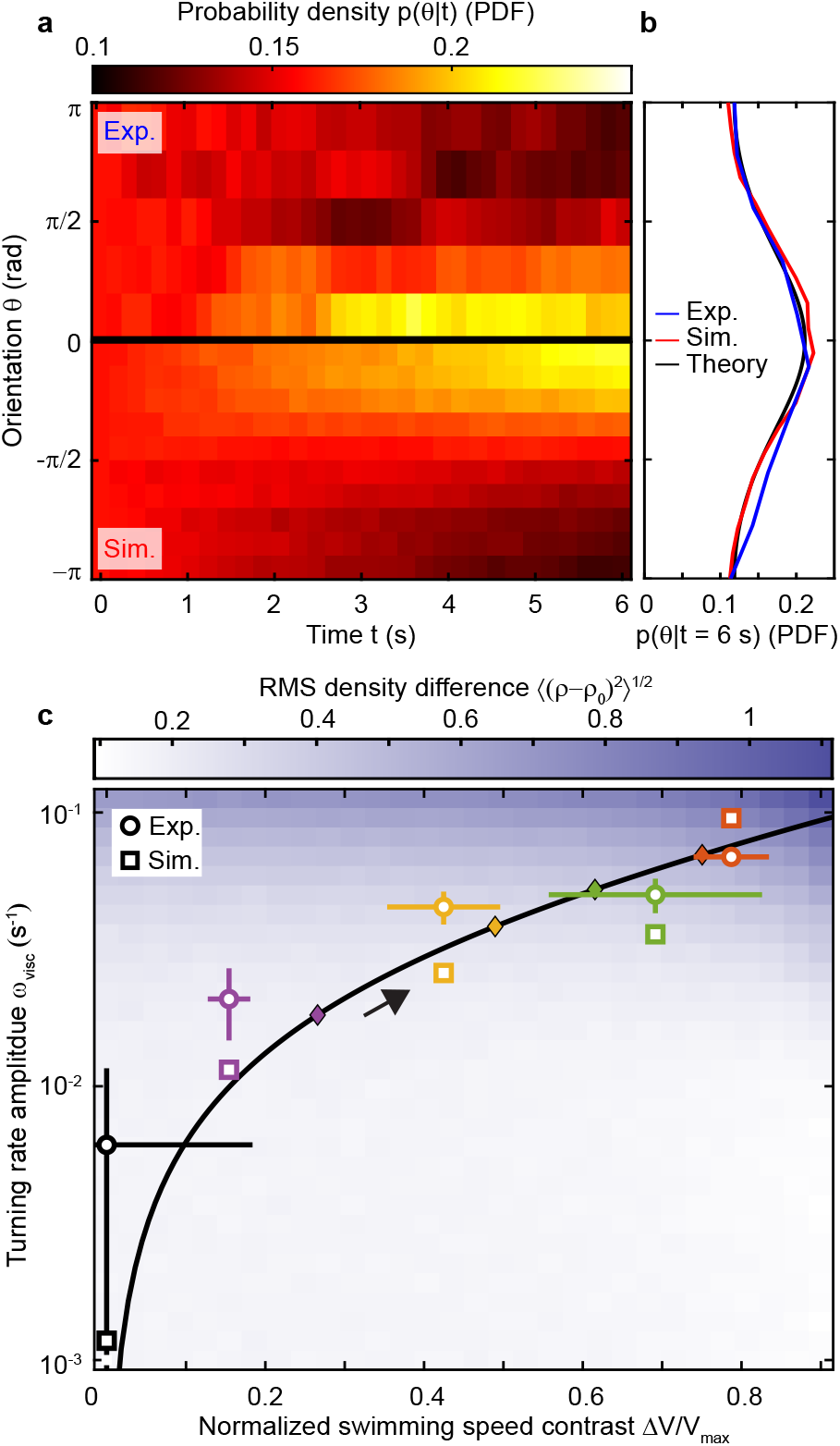
Simulations illustrate the competition between viscophobic drift and viscous slowdown driving cell accumulation. **a**, Time evolution of conditional probability density of cell orientation (▽*η* = 7.2 × 10^−3^ cP·*μ*m^−1^) for experiment (top) and Langevin simulation (bottom), showing condensation of cell alignment down the viscosity gradient (Extended Data Fig. 2). Experimental cell tracks selected from 400 ≤ *x* ≤ 600 *μ*m. **b**, Orientation distribution at t = 6 s from experiments and Langevin simulations (**a**) agree with a Fokker-Planck model (Supplementary Section 5.3). **c**, Root-mean-square (RMS) difference (colormap; see Methods) of cell density between Langevin simulations, *ρ*, and theory without viscophobic turning, *ρ*_0_ (Supplementary Section 4.1 and Extended Data Fig. 3). Squares are best match density profile simulations with experiments (circles; Fig. 1i). For circles, horizontal and vertical error bars propagated from Fig. 1h and Fig. 2d, respectively. Black curve is empirical scaling law (Supplementary Section 5.2), arrow indicates increasing ▽*η*, and diamonds correspond to experimental conditions (Fig. 1).

To elucidate the relative contributions of viscophobic turning versus viscous slowdown to cell transport, we employ a simplified, two-dimensional Langevin model. The simulated cell orientational dynamics are set by equation (1), and the translational dynamics are given by 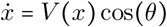 and 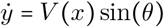 (Supplementary Section 5.1). The essential features of the viscosity dependence are captured by a spatially varying (linear) cell swimming speed profile, *V*(*x*), which is parameterized by the speed difference across the channel normalized by the maximal speed in the low viscosity zone, Δ*V*/*V*_max_. Resulting Langevin simulations (Fig. 1i, dashed colored lines) that best match the experimental cell density profiles faithfully reproduce the measured viscophobic turning rate (Fig. 3c, squares). The simulations also agree well with an empirical scaling law (Fig. 3c, black curve; Supplementary Section 5.2) based on the viscosity dependent swimming speed (Fig. 1f) and turning rate (Fig. 2d).

While viscophobic reorientation (Fig. 2d) and viscous slowdown (Fig. 1h) are intrinsically linked in experiments through the viscosity profile, simulations allow us to untangle these effects by independently varying *ω*_visc_ and Δ*V*/*V*_max_. Density profiles from Langevin simulations (Fig. 1i, dashed), *ρ*(*x*), and from theory based solely on the viscous slowdown, *ρ*_0_(*x*), are compared by examining the root-mean-square (RMS) density difference, 〈(*ρ* − *ρ*_0_)^2^〉^1/2^ (Fig. 3c, colormap; Extended Data Fig. 3). The sharp increase in RMS density difference with increasing viscophobic reorientation rate, *ω*_visc_, is indicative of the progressive skewing of the cell density toward the low viscosity region (Extended Data Fig. 3).

For a fixed *ω*_visc_, the viscous slowdown causes cell accumulation on the viscous side, yet viscophobic motility moderates this trend more efficiently as the swimming speed contrast is increased. In real systems, the simultaneous increase of *ω*_visc_ and Δ*V*/*V*_max_ (Fig. 3c, black arrow) indicates that viscophobicity dominates for large viscosity gradients. The growing influence of viscophobic motility with increasing gradient presents an ecologically valuable asset for the escape or repulsion of cells from viscous environments^1,2,5^.

Our work illustrates an important transport mechanism for microalgae stemming from the competition between viscophobic turning and viscous slowdown, which we anticipate will have a rich dependency on organism flagellation. The short flagellar mutant experiments and the three point force model show that negative viscotaxis is associated with the dominance of a flagellar thrust imbalance over a cell body drag imbalance in the gradient. Extending the model suggests that swimmers whose flagellar appendages are directly in line with the cell body – including many bacteria such as those probed in early experimental work^6,7^ – would exhibit positive viscotactic turning. This prediction remains to be experimentally demonstrated, but such positive viscotactic turning would exhibit a cooperative effect with viscous slowdown and exacerbate swimmer accumulation in high viscosity. While the observed viscophobic turning kinematics appear highly consistent with the proposed hydrodynamic mechanism, our work does not entirely rule out the potential for mechanosensing. Complementary studies at the level of single flagella will also be required to fully elucidate the potential impact of viscosity gradients on flagellar beat patterns and coordination^13,33,34^. The demonstration of viscophobic turning has the potential to refine our understanding of cell motility in complex microbiological systems, where the impact of viscosity gradients on cell motility have been widely overlooked, ranging from biofilms^4^ to protective mucus layers^2,3^ to endangered coral ecosystems^1^. The experimental and analytical approaches established in this work will pave the way for understanding the manifestation of viscophobic motility in a range of important microorganisms and in complex fluid rheologies.

## Methods

### Cell culturing and media preparation

*Chlamydomonas reinhardtii* (www.chlamycollection.org) wild-type (CC-1690 wild type mt+, WT) and short-flagella mutants (CC-2347 shf1-277 mt−) were grown in 100 mL of minimal media (M1)^40^ at 22°C using gentle aeration and synchronized on a 14 hr on and 10 hr off light cycle. Cells were subcultured (10 mL) every three days at mid-exponential phase. Viscous media was prepared with polyethylene oxide (PEO; molecular weight, 400,000 g/mol; Sigma-Aldrich) in M1 (Extended Data Fig. 10) at PEO concentrations of 0%, 0.05%, 0.1%, 0.25%, 0.5%, and 0.82% (w/v), which remain Newtonian (overlap concentration, c* ≈ 3% (w/v))^41,42^. PEO solutions were mixed for 2 hrs at 40 rpm (tube rotator; Fisherbrand), and 0.2% (w/v) bovine serum albumen was added to all solutions to reduce sticking to microchannel walls.

### Microfluidics

Microfluidic devices were made using standard soft lithography techniques^43^. Polydimethylsiloxane (PDMS; Dow Corning SYLGARD 184) channels were cast on photoresist (Microchem) molds, fabricated via photolithography, and plasma bonded to standard glass slides. Circular observation channels (diameter, 5 mm; height, 250 *μ*m) were used for bulk cell motility and microrheology measurements. Viscosity gradient generation channels (Fig. 1b) were designed with three inlets (width, 0.5 mm) carrying a viscous PEO solution, cell suspension, and M1 media, respectively. The three solutions were flow stratified for a minimum of 2 min using two syringe pumps (Harvard Apparatus), where flow rates were adjusted to maintain a 4:1:4 ratio of the stream widths (Supplementary Section 1). Upon halting the flow, a monotonic viscosity profile, *η*(*x*), slowly evolves over 90 min (Extended Data Fig. 1) due to the relatively small PEO diffusivity^44^. The thin central band of *C. reinhardtii*, bracketed by fresh M1, ensured no chemotactic bias was introduced^27^ (Supplementary Section 3.2).

### Microrheology and viscosity profile measurement

The viscosity of bulk Newtonian PEO solutions was measured through well-developed microrheology approaches^45^ by seeding with dilute (0.5 *μ*L/mL) tracer particles (radius, *R* = 0.25 *μ*m; 2% solid; Life Technologies). Tracers were imaged using epifluorescence microscopy (10× 0.5NA objective; Nikon Ti-e) and recorded (Blackfly S, FLIR Systems) at 4 fps for 25 s. Mean square displacements (MSD) of 488-1,153 particle trajectories, 〈Δ*x*^2^〉, were measured over time, Δ*t*, via particle tracking. A linear fit (Extended Data Fig. 4) of the MSD, 〈Δ*x*^2^〉 = 2*D*Δ*t*, yielded the diffusivity, *D*, which was used to determine the viscosity from the Stokes-Einstein relation, *D* = *k*_B_*T*/6*πηR* (Boltzmann constant, *k*_B_; absolute temperature, *T*). Measured bulk viscosities (Extended Data Fig. 10) compared well with previously reported results^41^. Spatially-resolved viscosity profiles, *η*(*x*), generated by the viscosity gradient assay (Fig. 1g), were measured using an identical approach to bulk experiments, but particle tracks were partitioned into 11 equally spaced bins across the channel width (Extended Data Fig. 5). Time evolution of the viscosity profile was evaluated by acquiring video of fluorescent tracer motion (10× 0.2NA objective) for 250 frames (63 s long) every 10 minutes for 90 min (Zyla, Andor Technology). The local viscosity was measured from the MSD (Extended Data Fig. 5) in each spatially resolved bin over the 90 min experiment. Transient variations in viscosity rapidly decreased after 30 min (Extended Data Fig. 1). Thus, we analyze our primary quasi-steady viscotaxis assay results from 30-90 min after halting the flow, and we report the average viscosity profile over this time window (Fig. 1g). Viscosity gradient, ▽*η*, is calculated from a linear fit to the viscosity profile in the middle third of the microchannel. The viscosity gradient for each viscotaxis experiment is the average gradient within the 30-90 min interval (Extended Data Fig. 8). See Supplementary Section 2 for details.

### Cell imaging and tracking

Imaging for viscotaxis experiments with *C. reinhardtii* was performed using phase-contrast microscopy (10× 0.2NA objective; Nikon Ti-e). To ensure that phototactic effects did not bias cell motility measurements^29^, experiments were performed in a darkened room equipped with black-out shades, a light shield was constructed to block stray light, and the microscope’s transmitted light source was equipped with a 665 nm longpass filter (Thorlabs, Inc.), which was outside of the spectral detection range of *C. reinhardtii* ^46^. Control experiments in the absence of viscosity gradients (Extended Data Fig. 6) confirmed that no phototactic bias was present. Upon halting the flow and initializing the viscosity gradient formation, videos were recorded (Zyla, Andor Technology) for 1000 frames each at 30 fps, every 5 min for the first 90 min. Dilute cell suspensions (≈ 66 cells per frame) ensured high fidelity cell tracking and negligible cell-cell interactions^47^ and mixing. Cells were tracked using a custom predictive particle tracking code written in MATLAB (MathWorks). Briefly, cell centroids were identified by intensity thresholding^48,49^ and their trajectories were reconstructed via distance minimization of quadratic projections of cell motion in comparison to new candidate cell locations^50^. Three to four replicate experiments were performed for each viscosity gradient for the wild-type experiments and one to three replicates for the short-flagella mutant experiments. Spatial cell density and swimming speed profiles were measured by partitioning the channel into 11 equally spaced bins (≈ 90 *μ*m wide; Fig 1h,i). To quantify the cell turning rate, *ω*, as a function of orientation (Fig. 2), trajectories were smoothed with a Gaussian filter (1 s standard deviation) to remove high frequency fluctuations, stemming from the natural helical trajectories of *C. reinhardtii*^51^. Occasional stuck or non-motile cells were removed from data by requiring a minimum ratio of track contour length to radius of gyration ≥ 4 and 6 for the wild-type and short mutant, respectively.

### Rotational cell diffusion

Rotational diffusion of (wild-type) *C. reinhardtii* was measured by recording cells (2× 0.1NA objective; Blackfly S, FLIR Systems) for 10 min at 15 fps and 10 fps in bulk 0.996 cP and cP media, respectively. Cell tracks with a minimum duration of 15 s were filtered as described above, and a minimum mean cell swimming speed of 54.2 *μ*m/s and 15.0 *μ*m/s was required for the 0.996 cP and 4.48 cP media, respectively. The MSDs (Extended Data Fig. 7) were fitted (Extended Data Fig. 7, solid lines) using the analytical result^52^,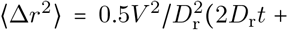 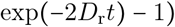, where the two-dimensional (2D) projections of the cells’ 3D random walks accurately capture the amplitude of their effective in-plane rotational noise^53^. For measured mean swimming speeds *V* = 74.7 *μ*m/s and 22.7 *μ*m/s, the resulting rotational diffusion coefficients were *D*_r_ = 0.070 s^−1^ and 0.068 s^−1^ for 0.996 cP and 4.48 cP, respectively.

### Langevin simulation parameters

Langevin simulations (Fig. 3 and Extended Data Fig. 2–3) were performed by numerically integrating the equations of motion for cell position and orientation using a fourth-order Runge-Kutta method with a time step of 0.001 s (Supplementary Section 5.1). Swimmers are uniformly seeded across the domain with random orientation, including: 500 swimmers for Fig. 3c and Extended Data Fig. 3 and 100,000 for Fig. 3a,b and Extended Data Fig. 2, which were integrated for 1000 s and 6 s, respectively. Based on measured cell motility parameters (Fig. 1–2), viscophobic rotation rate and viscous slowdown were independently varied via *ω*_visc_ and Δ*V*/*V*_max_, respectively. Typical, constant values were chosen for all simulations (Fig. 3 and Extended Data Figs. 2–3), including *V*_max_ = 100 *μ*m/s and D_r_ = 0.069 s^−1^. 2D Langevin simulations using *D*_r_ measured from 2D projections of cell trajectories (see above) have been shown to accurately reproduce measured cell transport properties^53,54^. Cell scattering from microchannel side walls was quantified^31^ from control experiments (▽*η* = 0 cP·*μ*m^−1^) and implemented as an empirical boundary condition (Supplementary Section 5.1). 900 total simulations spanning Δ*V*/*V*_max_ and *ω*_visc_ were performed for Fig. 3c. For a fixed Δ*V*/*V*_max_ (corresponding to a ▽*η*), the minimum root-mean-square (RMS) difference between experimental (Fig. 1i, solid lines) and simulation (Extended Data Fig. 3) density profiles determined the best match *ω*_visc_ from simulations (Fig. 3c, squares), which compared well with measured values (Fig. 3c, circles).

## Data availability

The data generated during the current study are available from the corresponding author upon reasonable request.

## Code availability

The algorithms and simulation codes are described in the Methods and Supplementary Information.

## Acknowledgements

We thank G.J. Elfring for helpful discussions and R.J. Henshaw for comments on the manuscript. This work was funded by NSF Awards CAREER-1554095 and CBET-1701392 to J.S.G.

## Author contributions

M.R.S and N.W. contributed equally to this work. M.R.S, N.W., and J.S.G. designed research; M.R.S. and N.W. performed experiments; M.R.S., N.W., and D.M.W. analyzed experimental data; N.W., D.M.W., M.R.S, and J.S.G. contributed to simulations and theoretical analyses; M.R.S., N.W., M.W., and J.S.G. wrote the paper.

## Additional information

Correspondence and requests for materials should be addressed to.

---

**Extended Data Fig. 1:**
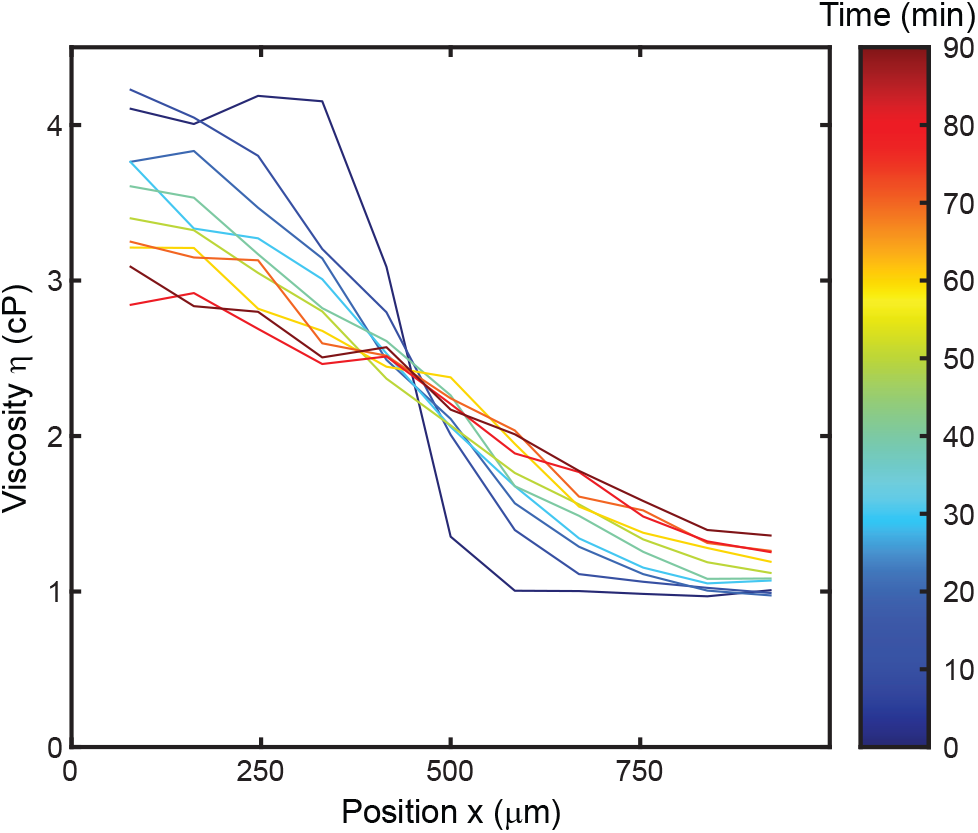
Time evolution of the viscosity profile. Transient diffusion of the PEO molecules results in a viscosity profile that evolves from a near step function to a smooth monotonic profile. Swimming cell assays are analyzed in the time range 30 ≤ *t* ≤ 90 min, when the gradient evolves more slowly, and the profile is approximately linear in the center of the channel. The slope of the viscosity profile is quantified by fitting a line to the five central points in the channel. The reported viscosity gradient, ▽*η* is the average slope of the viscosity profiles from 30-90 min (Extended Data Fig. 8).

**Extended Data Fig. 2:**
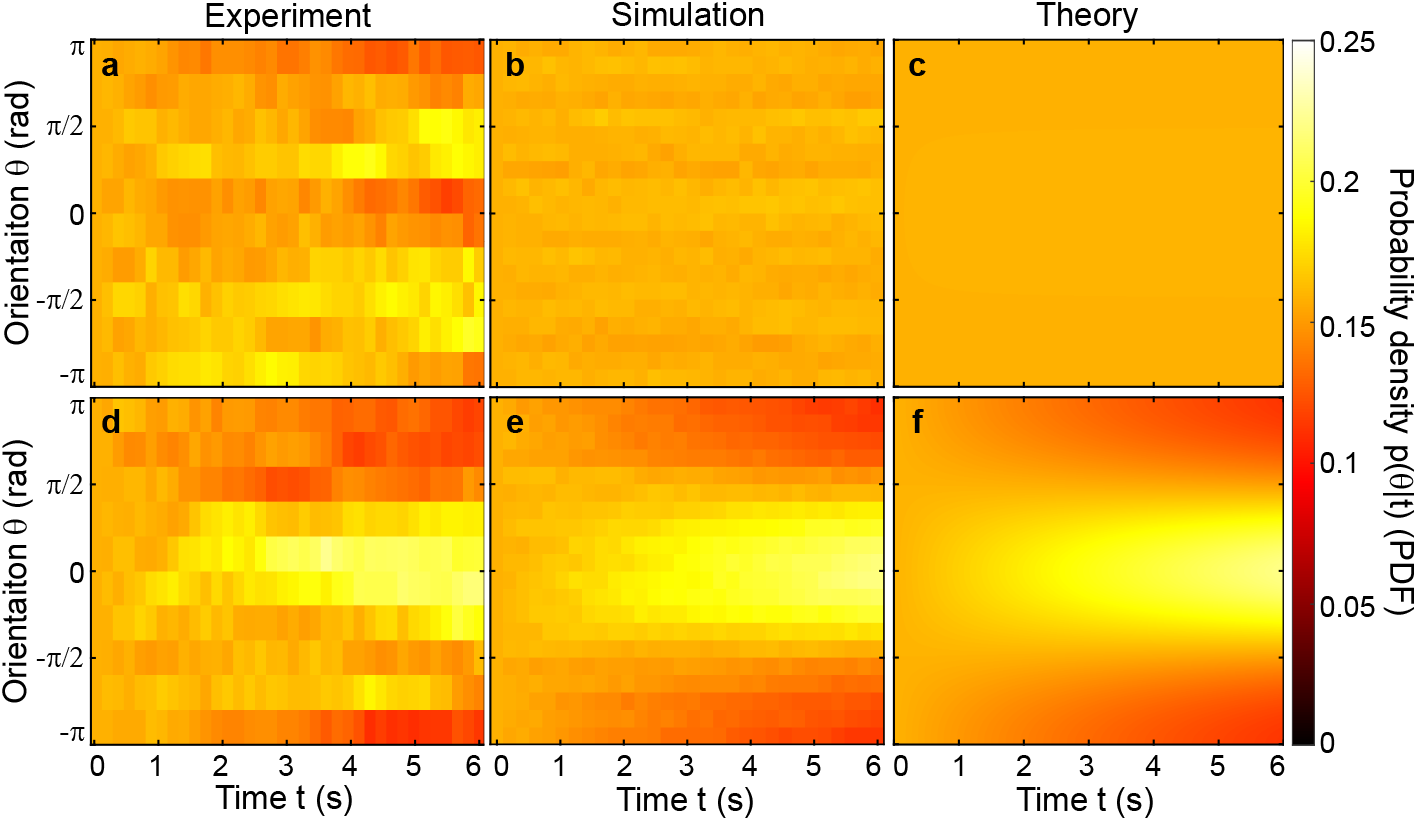
Time evolution of swimming cell orientation distribution in a viscosity gradient. **a-c**, In the absence of a viscosity gradient or viscophobic turning, an initially random uniform distribution of cell swimming orientations remains random, as illustrated across experiments, Langevin simulations, and a Fokker-Plank model, respectively. **d-f**, In contrast, in the presence of a viscosity gradient or orientation dependent viscophobic turning rate, the orientation of cells readily condenses in the down-gradient direction. Experimentally measured cell swimming trajectories were sub-sampled from the central portion (400 *μ*m ≤ *x* ≤ 600 *μ*m) of the microfluidic gradient generating channel with a uniform orientation distribution *p*(*θ*|*t*) at *t* = 0 for ▽*η* = 0 cP·*μ*m^−1^ (**a**; control, ≈ 1,170 trajectories) and for ▽*η* = 7.2 × 10^−3^ cP·*μ*m^−1^ (**d**; ≈ 1,330 trajectories). Peaks in **a** at ≈ 6 s are due to cell collisions with microchannel walls. Langevin simulations in the absence (**b**) and presence (**e**) of viscophobic turning (*ω*_visc_ = 0.07 s^−1^, Δ*V*/*V*_max_ = 0.776) show quantitative agreement with experiments having matching conditions (Extended Data Fig. 3 and Fig. 3a,b) in **a** and **d**, respectively. A one-dimensional Fokker-Planck model of the cell swimming orientation distribution (Supplementary Section 5.3) likewise captures the time evolution of the conditional probability density for the control (**c**) and corresponding maximum viscosity gradient conditions (**f**; Fig. 3b).

**Extended Data Fig. 3:**
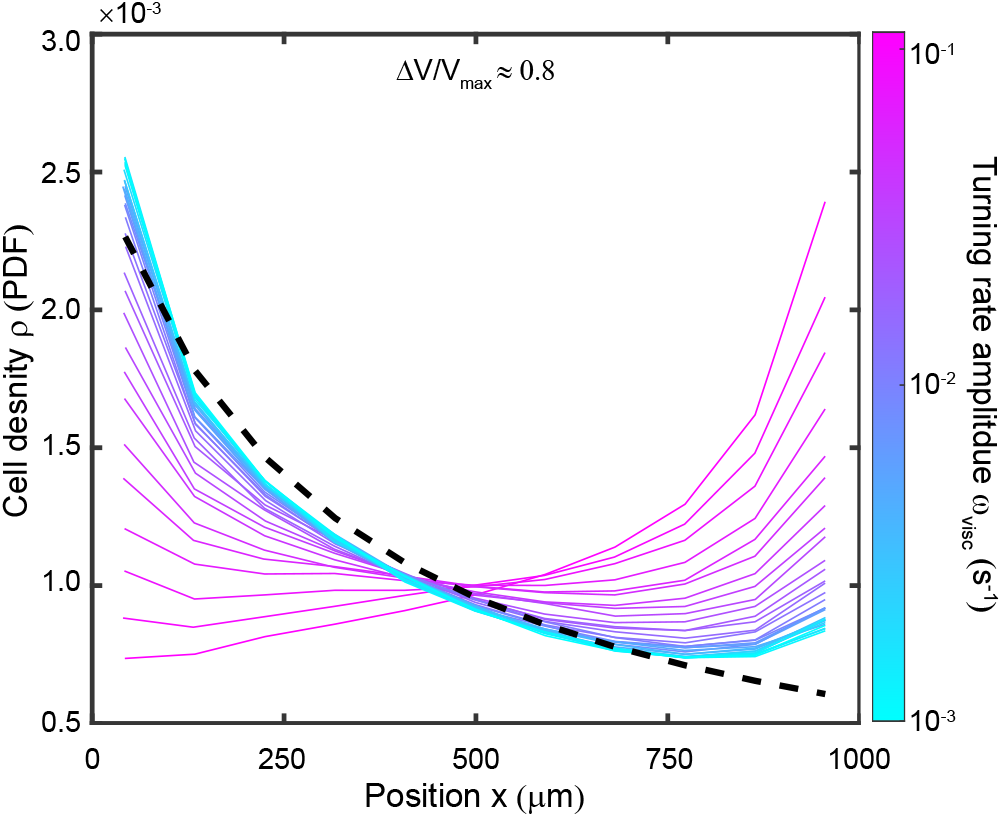
Comparison of cell density profile due to (viscous) slowdown and Langevin simulations incorporating viscophobic turning. Langevin simulations enable independent exploration of the parameters Δ*V*/*V*_max_ and *ω*_visc_, and include an empirical wall scattering boundary condition (Supplementary Section 5.1). Cell density profiles, *ρ*(*x*), are shown for fixed Δ*V*/*V*_max_ and a range of turning rate amplitudes (solid lines). The resulting density profiles are compared to a theoretical distribution, *ρ*_0_, for cells with spatially varying swimming speed in the absence of viscophobic turning (black dashed curve; Supplementary Section 4.1).

**Extended Data Fig. 4:**
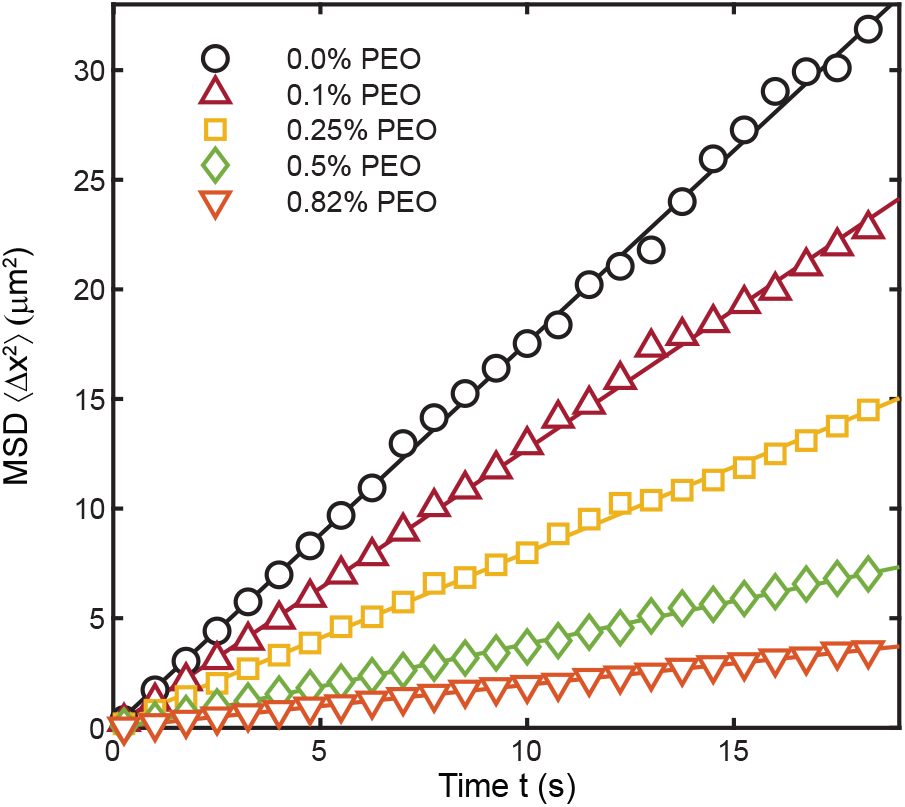
Mean square displacements of tracer particles for viscosity measurements in bulk fluid. Microrheology quantifies the viscosity of PEO solutions in M1 media at various concentrations. The slopes of linear fits (solid lines) to the mean square displacements (MSDs; markers) yield the tracer particle diffusion coefficients, which are used to determine the viscosity of the solution via the Stokes-Einstein relation (Supplementary Section 2.1). Measured viscosities are listed in Extended Data Fig. 10.

**Extended Data Fig. 5:**
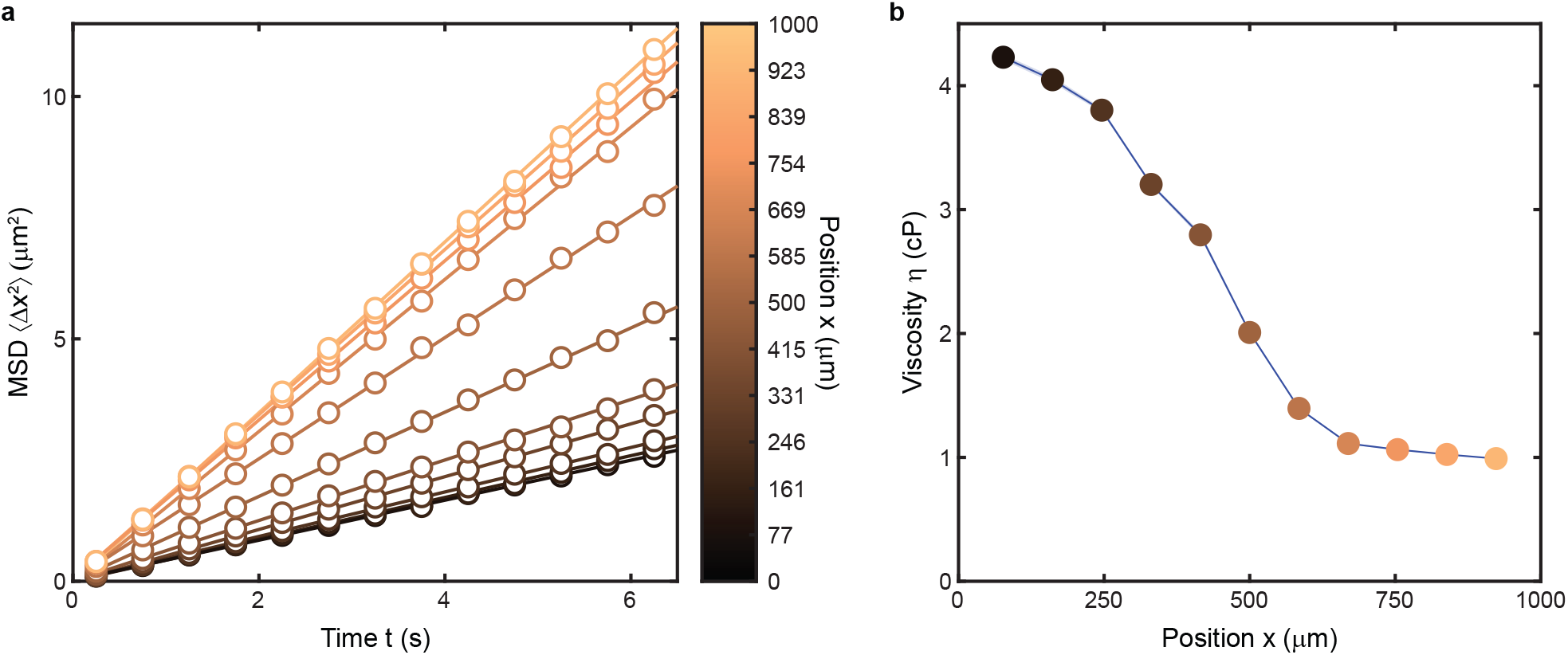
Spatially resolved microrheology quantifies the viscosity profile in a microfluidic device. **a**, The mean square displacement (MSD) of tracer particles in different locations across the microfluidic viscosity gradient device at time, *t* = 10 min (▽*η* = 3.4 × 10^−3^ cP·*μ*m^−1^; see also Extended Data Fig. 1). Solid lines are linear fits to the MSDs. **b**, The slopes of the tracer particle MSDs in **a** are used to determine the spatially resolved viscosity profile, *η*(*x*), across 11 bins of the microchannel width using Stokes-Einstein relation (Supplementary Section 2.2). Shaded region (smaller than the markers) represents the propagation of uncertainty from the error of the fit in **a**.

**Extended Data Fig. 6:**
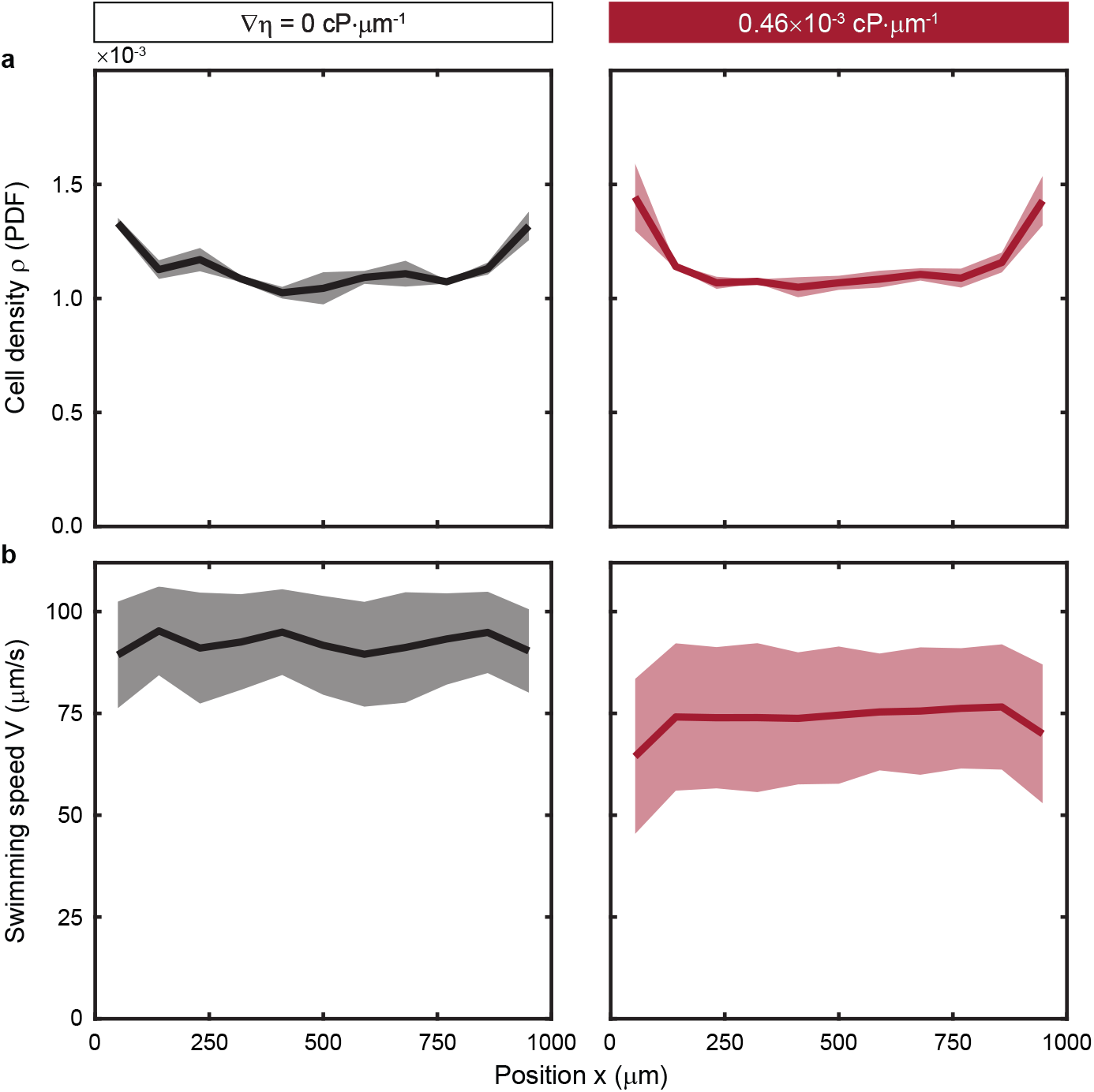
Control experiments show negligible chemotaxis and chemokinetic effects for wild-type *C. reinhardtii* in PEO gradients. **a,b**, Viscotaxis assays reveal the local, measured cell density (**a**) and mean swimming speed (**b**) remain spatially uniform for small viscosity gradients. Base PEO concentrations of 0.05% in “Media + PEO” inlet (▽*η* = 0.46 × 10^−3^ cP·*μ*m^−1^; N=3; right) are comparable to the control experiment with no viscosity gradient (▽*η* = 0 cP·*μ*m^−1^; N=4; left, repeated from Fig. 1h,i). For these control experiments, the base PEO concentration (0.05%) is nearly the same order of magnitude as the maximum gradient (0.82%), yet has no appreciable effect on the local viscosity. Thus, the lack of discernible cell accumulation and the lack of statistically significant spatial variations in swimming speed indicate that PEO likely does not act as a chemoattractant nor does it have any chemokinetic effect on *C. reinhardtii*. The shaded areas represent the standard error.

**Extended Data Fig. 7:**
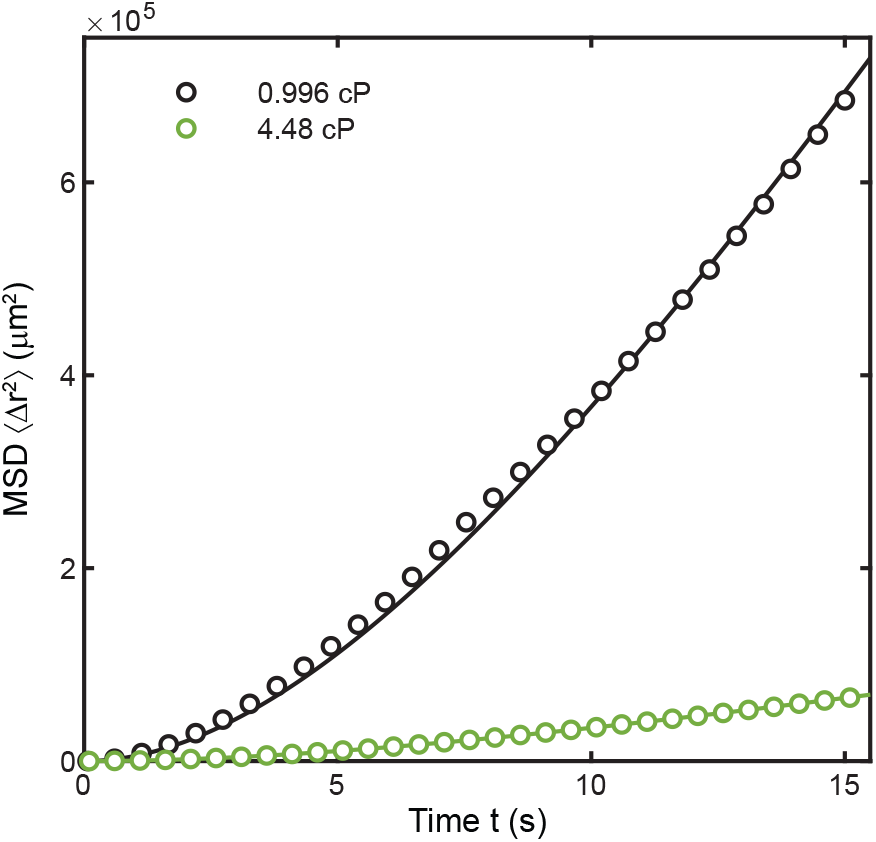
Measurement of mean square displacement (MSD) of wild-type *C. reinhardtii* in uniform viscosity to determine rotational diffusion coefficient. Mean square displacements of *C. reinhardtii* were measured in 0.996 cP (≈ 1,800 trajectories) and 4.48 cP (≈ 10,700 trajectories) bulk viscosities. The resulting MSDs were fitted using an analytical result for a persistent random walk^52,53^, 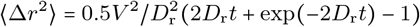, where the ensemble averaged cell swimming speeds, *V*, are known independently (74.7 *μ*m/s and 22.7 *μ*m/s, respectively), and the rotational diffusion coefficient, *D*_r_, is the lone fitting parameter. The resulting rotational diffusion coefficients were *D_r_* = 0.070 s^−1^ and 0.068 s^−1^. As a sensitivity analysis, varying the fit window from 10 s to 20 s resulted in *D_r_* = 0.071 ± 0.006 s^−1^ and 0.068 ± 0.001 s^−1^ for 0.996 cP and 4.48 cP, respectively, where the error is the standard deviation of the resulting rotational diffusion coefficients.

**Extended Data Fig. 8:**
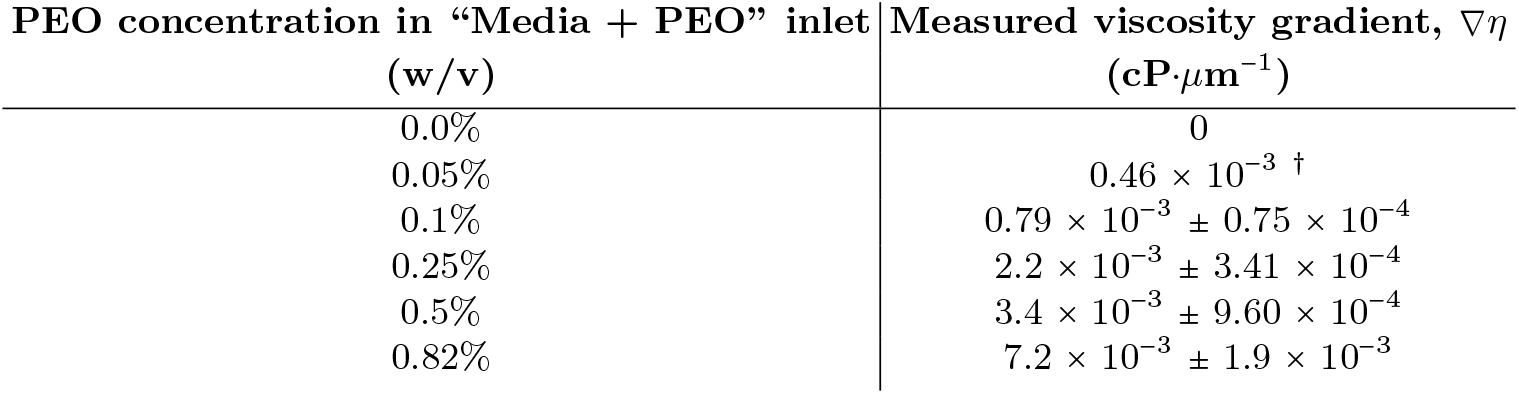
Measured viscosity gradients for viscotaxis experiments. Viscosity gradients, ▽*η*, for each of the tested gradient conditions were generated by controlling the bulk PEO concentration/viscosity in the “Media + PEO” inlet (Fig. 1b). Viscosity gradients were measured by fitting a line to the measured viscosity profiles in the central third of the channel. The slopes of the fits were then averaged to determine the mean ▽*η* over the experiment time window of 30-90 min. The ^†^ indicates interpolation by quadratic fit. Uncertainty represents the standard deviation of the fitted viscosity profile slopes from 30-90 min (Extended Data Fig. 1).

**Extended Data Fig. 9:**
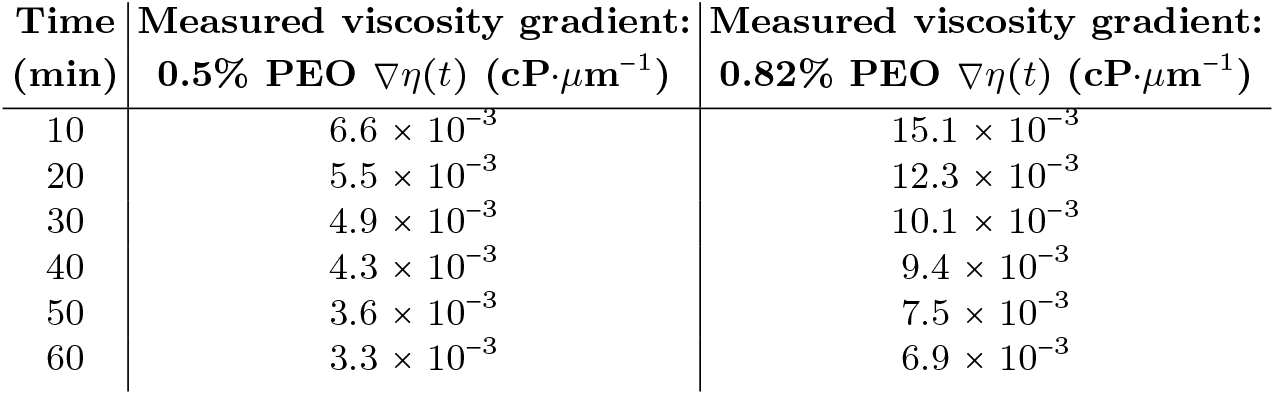
Measured transient viscosity gradient for viscotaxis experiments. Viscosity gradients were measured by fitting a line to the measured viscosity profiles in the central third of the channel. The slopes of the fits determine the unsteady viscosity gradient, ▽*η*(*t*), at early times in the assay as the PEO molecules diffuse across the microfluidic device. 0.5% and 0.82% PEO transient viscosity gradient data correspond to steady experiments (30-90 min post flow) for ▽*η* = 3.4 × 10^−3^ cP·*μ*m^−1^ and ▽*η* = 7.2 × 10^−3^ cP·*μ*m^−1^ (green and orange in Fig. 1), respectively.

**Extended Data Fig. 10:**
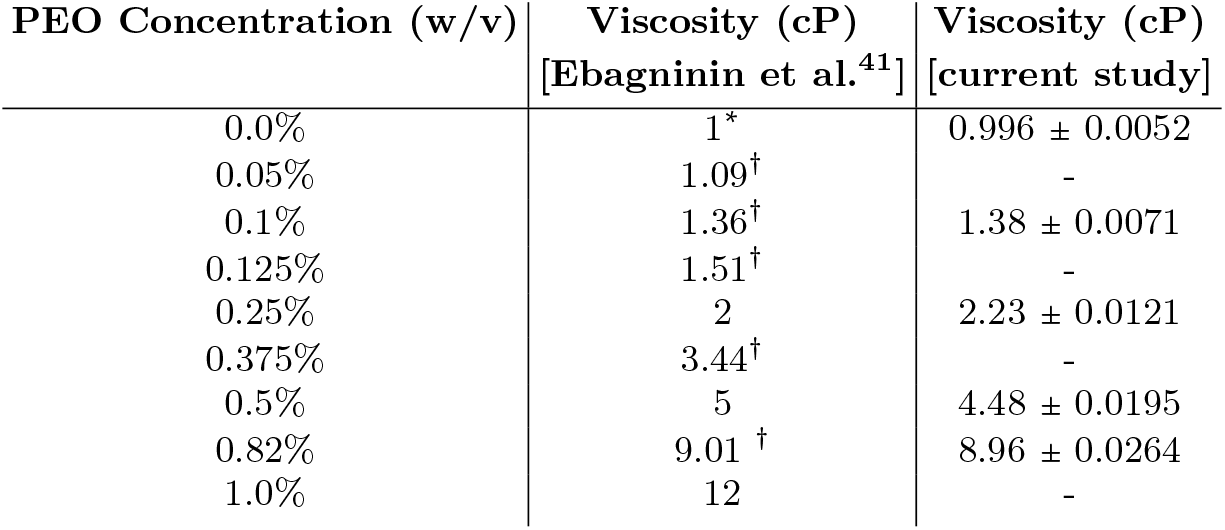
Viscosity of bulk PEO solutions. The dynamic viscosities of aqueous PEO solutions of various concentrations reported in the literature^41^ compared to the present study in M1 media (measured by microrheology; ≈ 22°C). The * indicates the known viscosity value at 20°C^55^ and the ^†^ indicates a quadratic interpolation of Ebagninin et al.^41^. Experimental uncertainty represents the propagation of error from the calculation of the diffusion coefficient.

## Supplementary Information

### 1. MICROFLUIDIC VISCOSITY GRADIENT ASSAY

Monotonic viscosity profiles of various gradient magnitude were generated by prescribing the viscosity of the solutions in each of the three inlet channels (Fig. 1b and Fig. S1): (*i*) The leftmost inlet contained a viscous polyethylene oxide (PEO) solution (molecular weight (MW), 400,000 g/mol; Sigma-Aldrich) based in minimal (M1) media (solution, 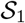; Extended Data Fig. 10) [1]. (*ii*) The center channel contained a dilute *C. reinhardtii* suspension in M1 media (solution, 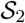). (*iii*) The rightmost channel contained M1 media (solution, 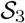). In the case of viscosity profile measurements, the *C. reinhardtii* suspension in 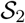 was replaced by M1 media and all three solutions were seeded with a dilute suspension of tracer particles (see §2). To initiate the viscosity gradient, two syringe pumps (one for 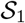 and one for 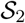 and 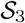; Harvard Apparatus) were flowed at a high rate of speed for 2 min to establish stratification between the three viscous solutions in the main test channel (Fig. S1). At the start of the assay, both syringe pumps were abruptly stopped, and videos of cell or tracer particle motion were captured periodically over the course of the experiment. Upon halting the flow, the initially step-wise concentration profile (and consequently viscosity profile, *η*(*x*)) of PEO slowly evolved over the course of 90 min into a smooth concentration/viscosity profile (Extended Data Fig. 1) due to the small diffusivity (*D* ≈ 2 × 10^−7^ cm^2^/s) of the relatively large PEO molecules [2].

We chose the target widths of the initial fluid bands for 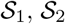, and 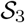 in the test section to be *w*_1_ = 4W/9, *w*_2_ = *W*/9, and *w*_3_ = 4*W*/9, respectively (Fig. S1). The thin central band of *C. reinhardtii* in M1 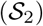 ensured that no bias is introduced in the assay due to the starting position of the cells. Furthermore, bracketing the cell suspension by solutions based on fresh M1 media (i.e. 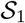 and 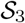) ensured that there were no extraneous chemical gradients that might bias motility, for example due to the consumption of resources by the cells [3]. Owing to the differential viscosities, *η*_*i*_, of the PEO and M1 solutions, initial flow rates, *Q*_*i*_, were adjusted to maintain the symmetry of the desired band widths, *w*_*i*_, established above for the main channel for 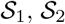, and 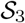. From the predicted hydraulic flow resistance of each solution in the main channel [4], the desired band widths are related to the respective flow rates and viscosities of the solutions approximately by *w*_*i*_*η*_*i*_/*Q*_*i*_ = constant. Due to the dilute nature of the cell suspensions, we assumed that the viscosities of 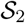 and 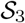 are the same, *η*_2_ = *η*_3_, as the M1 media alone [5]. Additionally for viscosity profile measurements, we assumed that the addition of dilute tracer particles (see §2.1) does not augment the viscosities of the respective solutions. Solutions 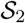 and 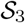 were flowed through the device using glass syringes (Hamilton) with various barrel radii, *r_i_* (*r*_3_ > *r*_2_), and controlled by a single syringe pump resulting in relative flow rates *Q*_3_ = 17 *μ*L/min and *Q*_2_ = *Q*_3_(*r*_2_/*r*_3_)^2^ = 4.25 *μ*L/min. This choice ensured achieving the prescribed flow stream widths. The viscous PEO solution, 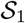, was injected using a second, separate syringe pump and syringe with an initial flow rate *Q*_1_ = *Q*_3_*η*_3_/*η*_1_. Phase contrast microscopy enabled visualization of the stratified fluid interfaces, and the flow rate *Q*_1_ was fine tuned to ensure that the initial, desired symmetry of the stratified solutions (*w*_1_ = *w*_3_) was maintained for all experiments. The final adjusted flow rates were *Q*_1_ ≈ 17, 14, 11, 7, 5 and 3.5 *μ*L/min for PEO solutions with concentrations 0%, 0.05%, 0.1%, 0.25%, 0.5%, and 0.82% (w/v), respectively (Extended Data Fig. 10). Finally, during control experiments, a negligible change in swimming speed (Extended Data Fig. 6b, left) compared to the bulk measurements (Fig. 1f) verified that this assay had no negative effects on the motility of the robust *C. reinhardtii* cells.

### 2. MICROSCALE VISCOSITY MEASUREMENTS

Directly quantifying spatially varying viscosity on sub-millimeter scales was necessary to understand the viscotactic transport of swimming cells in viscosity gradients. To measure the viscosity of both the spatially varying viscotaxis experiments and bulk PEO solutions, we adapted techniques from well-developed microrheology approaches [6, 7].

#### 2.1. Bulk media

For uniform, bulk fluids, we measure the Brownian thermal motion of micron-sized tracer particles suspended in viscous fluids of interest [8], which we then relate to the viscosity, *η*. For simple, Newtonian fluids, the viscosity is quantified by measuring the mean square displacements (MSD), 〈Δ*x*^2^〉, of microparticles over time, Δ*t*, through particle tracking. We focus on particle displacements, Δ*x*, perpendicular to the direction of the flow in the microfluidic devices to circumvent any minor tracer particle drift due to residual flows in the channels during the assay. The diffusion coefficient, *D*, of the tracer particles is first measured from a linear fit to the MSD over Δ*t*:

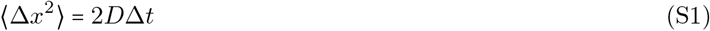

Applying the Stokes-Einstein relation, the viscosity, *η*, is inferred from the measured diffusion coefficient (equation (S1)) [6]:

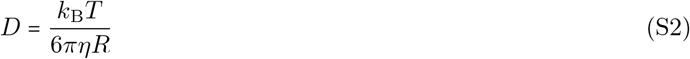

where *k_B_* is the Boltzmann constant, *T* = 295 K is the absolute ambient temperature, and *R* is the radius of the tracer particles.

To quantify the media viscosity, epifluorescence microscopy (Nikon Ti-e) captured the diffusive motion of fluorescent tracer particles. Viscous media – M1 media with PEO concentrations of 0.1%, 0.25%, 0.5%,and 0.82% (w/v) – was seeded with 0.5 *μ*L/mL of fluorescent tracer particles (*R* = 0.25 *μ*m radius; 2% solid; carboxylated FluoroSpheres, Life Technologies) and placed in a microfluidic observation chamber, where the inlet and outlet were sealed with wax. Using a 10x magnification objective (0.5NA) with an additional 1.5x magnification multiplier lens, videos of the particles were recorded (Blackfly S, FLIR Systems) at a frame rate of 4 fps for 25 s. Particle tracking (see Methods) was used to measure the 〈Δ*x*^2^〉 of 488-1,153 particle trajectories. With increasing concentrations of PEO, the diffusion coefficient (equation (S1)) decreases (Extended Data Fig. 4), revealing an increase in bulk viscosity. The experimentally measured bulk viscosities are reported in Extended Data Fig. 10, and compare well with previously reported results [9]. Furthermore, PEO suspensions are known to remain Newtonian under these concentration conditions (*c* < 1% w/v), which are well below the transition to semi-dilute solutions, *c*\* ≈ 3% (w/v) [9].

#### 2.2. Viscosity profile

Using the same microfluidic device and conditions as described in §1 and in the main text for the cell motility assays (Fig. 1b), the spatially varying viscosity profile in the channel test section was quantified, in the absence of *C. reinhardtii*, by seeding the media with the fluorescent tracer particles. Following the viscosity gradient assay procedure along with the bulk viscosity measurement procedure above, epifluorescence microscopy captured the diffusive motion of the tracer particles within the viscosity gradient. Viscosity profiles were quantified using a 10x magnification objective (0.2NA) with an additional 1.5× magnification multiplier lens and 10 videos were recorded at 4 fps for 250 frames (63 s long) every 10 minutes for 90 min (Zyla, Andor Technology). For the control experiments, in which solution 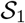 contained no PEO, videos were recorded with the same parameters as the bulk media at 4 fps for 100 frames every 3.5 minutes for 10 minutes (Blackfly S, FLIR Systems). By spatially resolving the seeded tracer particles using particle tracking, the channel was partitioned into 11 equally spaced bins, where 〈Δ*x*^2^〉 was calculated over time, Δt, for each bin (Extended Data Fig. 5). Following this method over the course of the experiment, the viscosity profile’s evolution over time was realized (Extended Data Fig. 1). After about 30 min, the strong transient variations in the viscosity profile decrease. Thus, we report and analyze our primary results 30 to 90 minutes after ceasing the initial stratifying flow, when the viscosity profile varies weakly in time. While the slope of the viscosity profile changes slightly from 30 min to 90 min, the motility of *C. reinhardtii* enables the cells to rapidly sample the channel, resulting in quasi-steady cell transport (see §3.3). This substantial difference in time evolution of the PEO concentration and cell concentration justifies the use of MW 400,000 g/mol PEO molecules for the generation of the viscosity profile. The average viscosity profile, over the 30-90 min, was calculated by averaging the local measured viscosity for each bin over time (Fig. 1g).

From each measured viscosity profile, *η*(*x*), the viscosity gradient, ▽*η*, was calculated from a linear fit to the middle third of the viscosity profile in the center of the microfluidic channel. The viscosity gradient reported for each experiment throughout the main text is the average gradient within the 30-90 min interval (Extended Data Fig. 8).

### 3. CONTROL EXPERIMENTS

Wild-type *C. reinhardtii* is a biflagellate (Fig. 1a) with a slightly prolate body (typical diameter of 10 *μ*m) [10]. These cells have two 10-12 *μ*m long anterior flagella (≈ 0.2-0.3 *μ*m diameter) [11, 12], and swim with a breast stroke pattern (≈ 50-60 Hz beat frequency) [10]. Short-flagella *C. reinhardtii* mutants (Methods), which are used in select experiments in the main text, are similar to wild-type cells except their flagella are 6-8 *μ*m long [13]. Below, we discuss a series of control experiments to complement those in the main text. Unless otherwise noted, experiments are performed with wild-type cells.

#### 3.1. Bulk viscosity swimming speed

In order to determine the effects of spatially inhomogeneous viscous media on the transport properties of wild-type *C. reinhardtii*, we first examined the effect of enhanced bulk viscosity on the motility of these cells by quantifying their ensemble average swimming speed, 〈*V*〉. The cells were gently resuspended in PEO solutions of various concentration (and thus viscosities) prepared in M1 media (Extended Data Fig. 10). To minimize dilution of the viscous media in the resuspension process, the cell suspension was washed three times: (*i*) 1 mL of cell suspension was pipetted into a 1.5 mL microcentrifuge tube and centrifuged (Eppendorf) at 2,800 rcf for 30 seconds to form a soft pellet, (*ii*) the supernatant (≈ 0.9 mL) was removed and replaced with an equal amount of viscous media, and (*iii*) the entire process was repeated two more times. The cells were pipetted into a PDMS microfluidic observation chamber and the inlet and outlet were sealed with wax to eliminate residual flow. Care was taken to ensure that the cells were not damaged during the pipetting and resuspension steps, and the robustness of motility was confirmed both visually and through quantitative video microscopy analysis (Fig. 1f).

Using video microscopy and particle tracking, the ensemble mean swimming speed, 〈*V*〉, of the cell trajectories over multiple videos were measured. For each bulk viscosity, 2-3 videos were recorded using the same specifications as for the viscotaxis experiments (see Methods). For bulk viscosities of *η* = 0.996 cP, 1.51 cP, 2.23 cP, 3.44 cP, 4.48 cP, and 8.96 cP (Extended Data Fig. 10), the ensemble averaged swimming speeds of *C. reinhardtii* were 〈*V*〉 = 92.2 *μ*m/s, 85.4 μm/s, 51.2 *μ*m/s, 30.3 *μ*m/s, 21.8 *μ*m/s, and 14.5 *μ*m/s, respectively (Fig. 1f). We observed a power law decrease of swimming speed with increasing viscosity of slope −0.934 (Fig. 1f, black line), which is consistent with prior observations [14] and constant thrust flagellar propulsion, 〈*V*〉 ~ *η*^−1^. Error bars in Fig. 1f represent the mean standard deviation across *N* = 3 videos (or the range in the case of *N* = 2 for 0.996 cP and 8.96 cP) for each bulk viscosity (each with ~ 2, 700 trajectories).

#### 3.2. Potential effects of chemotaxis and chemokinesis

Many microorganisms are capable of sensing, responding to, and metabolizing small polymers in suspension, which may introduce biases into experimental measurements [3, 15, 16]. Chemotaxis can cause cell accumulation in a chemical gradient [17] and chemokinesis can enhance cell swimming speeds in elevated chemical concentrations [18]. Through control experiments, performed at low concentrations of PEO in M1 media for wild-type *C. reinhardtii*, we demonstrate that neither chemotaxis nor chemokinesis affect the motility of *C. reinhardtii* in our system. The base PEO concentration (0.05% for solution 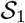; Fig. S1) was selected such that the viscosity was within 10% of the M1 media (corresponding to ▽*η* = 0.46 × 10^−3^ cP·*μ*m^−1^), while differing from the maximum concentration by approximately one order of magnitude. In repeating these assays with this smaller viscosity gradient, no cell accumulation was discernible in the cell density profile (Extended Data Fig. 6a, right) compared to the no gradient control experiment (▽*η* = 0 cP·*μ*m^−1^; Extended Data Fig. 6a, left). This observation indicates that chemotaxis effects are likely negligible in this system. Furthermore, no spatial variations in swimming speed were discernible for the lower viscosity gradient (▽*η* = 0.46 × 10^−3^ cP·*μ*m^−1^; Extended Data Fig. 6b, right) compared to the control. We note that a lower mean swimming speed was observed for ▽*η* = 0.46 × 10^−3^ cP·*μ*m^−1^, but this is consistent with the slightly elevated viscosity, the natural variations in population swimming speed, and the experimental uncertainty. Thus, we likewise conclude that there are no discernible chemokinesis effects by PEO on *C. reinhardtii* that affect the outcomes of our study.

#### 3.3. Potential effects of transient viscosity gradients and swimmer density

Viscosity gradients were generated in the microfluidic device by the diffusion of PEO polymer (see §1), which is an inherently unsteady process. This effect is apparent in viscosity profile measurements (Extended Data Fig. 1), and its effect on viscotaxis experiments was minimized by restricting our analysis to the 30-90 min window, post gradient formation (i.e. after the strongest transient regime subsides). As a test of the sensitivity of our results, using wild-type *C. reinhardtii*, we examined the variation of the local cell density, *ρ*, and swimming speed, *V*, profiles between 5-30 min, 30-60 min and 60-90 min for the maximum viscosity gradient tested (▽*η* = 7.2 × 10^−3^ cP·*μ*m^−1^; Fig. S2). For the cell density and swimming speed, we observe a mean percent difference across the 30-60 min and 60-90 min time intervals (relative to the 30-90 min interval) of less than 2.33% and 6.08%, respectively. This variation is well within acceptable tolerances and consistent with the uncertainty in our experiments.

The characteristic time scale for establishing the swimmer density profile (< 3 min) is significantly faster than the time to establish the viscosity gradient (≈ 30 min), emphasizing that the swimmer distribution is approximately stationary during the time of the experiments (30-90 min). We consider the characteristic timescale, *T*, for swimming cells to disperse throughout the microfluidic chamber (width, *W* = 1 mm). As a conservative estimate, we use the measured wild-type *C. reinhardtii* transport properties in the highest viscosity conditions (9 cP) occurring across all of our experiments, which provides the largest possible time scale. Under these conditions, we directly measured a cell swimming speed of *V*_s_ = 14.5 *μ*m/s (Fig. 1f) and an effective rotational diffusion coefficient of *D*_r_ = 0.07 s^−1^ (Extended Data Fig. 7). From these results, we obtain an effective cell diffusion coefficient of 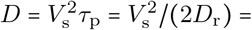 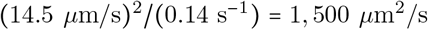. Because cells are injected into the center of the channel, the characteristic length scale for diffusion is *W*/2. Thus, the characteristic time for cell diffusion is *T* = (*W*/2)^2^/*D* = 166 s < 3 minutes. This analysis provides further evidence that transients due to swimmer motility have negligible impact on the swimmer density profiles (Fig. 1).

#### 3.4. Potential effects of diffusiophoresis

Concentration gradients of small dissolved molecules can cause a drift of suspended colloidal particles due to the Fickian solute flux, a process known as diffusiophoresis [19]. Through the control experiments below, we definitively confirm that diffusiophoretic effects, due to the established concentration gradients of PEO, have a negligible effect on the transport of *C. reinhardtii* in our viscotaxis experiments. In order to isolate the potential effects of diffusiophoresis, wild-type *C. reinhardtii* suspensions were rendered immotile while preserving the cell shape by fixing the cells in a 4% (v/v) solution of 37% formaldehyde (Sigma-Aldrich). Diffusiophoretic drift is known to scale with the solute concentration gradient. Therefore, control viscotaxis experiments were performed with fixed *C. reinhardtii* cells at the maximum tested viscosity (i.e. concentration) gradient of ▽*η* = 7.2 × 10^−3^ cP·*μ*m^−1^ under identical conditions and for the same 90 min duration as the experiments shown in the main text (Fig. S3a). The maximum displacement of the tracked non-motile cells in the viscosity (concentration) gradient direction was just Δ*x* ≈ 10 *μ*m (≈ 1 cell diameter or ≈ 1% of the channel width; Fig. S3b) during the 90 min experiment. The observed displacements are consistent with Brownian diffusion and are many orders of magnitude smaller than the effect of cell motility in the same time frame. This result illustrates unequivocally that diffusiophoresis does not affect the observed viscotactic/viscophobic transport of *C. reinhardtii* described in the main text (Fig. 1).

#### 3.5. Potential effects of swimming-induced mixing

Swimming cells including *Chlamydomonas* have been reported to enhance mixing, when in sufficiently high concentration (*ϕ* ~ 0.01 volume fraction) [20]. To minimize mixing effects, the cell concentration was maintained two orders of magnitude smaller at a volume fraction of *ϕ* ≈ 0.0002 (corresponding to ≈ 66 cells per video frame) for all experiments. The bulk diffusion enhancement due to a suspension of swimming *C. reinhardtii* was previously shown to grow linearly as *D*_eff_ = *D*_0_ + *αϕ* where *α* = 81.3 *μ*m^2^/s [20]. For the viscous polymer used in the present study (polyethylene oxide, PEO; diffusivity, *D*_0_ = 20 *μ*m^2^/s [2]), we expect a diffusion enhancement due to cell motility of just *αϕ*/*D*_0_ = 0.1% beyond the molecular diffusion of PEO, which is negligible. Our data (Fig. 1) also shows direct evidence that the viscosity gradient is essentially unaffected by the presence of the cells. We independently measured (*i*) the viscosity profile, *η*(*x*), using microrheology in the absence of cells (Fig. 1g), and (*ii*) the swimming speed dependence of the cells on bulk viscosity, *V*(*η*) (Fig. 1f). Together, these data enabled us to predict the spatial distribution of cell swimming speed in a viscosity gradient (Fig. 1h, solid gray curves), *V*(*η*(*x*)), which we compared directly to the measured cell swimming speed profile, *V*(*x*) (Fig. 1h, solid colored curves; see also §4.1). The agreement between these highly independent and repeatable measurements provide excellent confirmation that the cell motility does not significantly affect the viscosity gradient through mixing.

### 4. THEORETICAL CONSIDERATIONS

#### 4.1. Cell density based on measured bulk swimming speed

When cells are free from directional bias (e.g. viscophobic turning), local cell density, *ρ*_0_(*x*), is widely recognized to be inversely proportional to cell swimming speed [21], *V*(*x*), and in a one dimensional system:

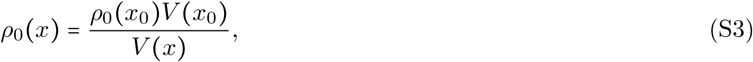

where *x*_0_ is an arbitrary reference location in the domain 0 ≤ *x* ≤ *W*. In the experiments shown in the main text, and generally in situations where speed *V*(*x*) is heterogeneous, we consider these predicted distributions *ρ*_0_(*x*) for unbiased random walks as the reference distributions because any deviation signals a bias in the motility. Here, we combine independent experimental measurements of cell swimming speed in bulk viscous media, locally measured viscosity profiles, and theory to predict these reference distributions, representing the expected cell density profiles in our system if there was no viscophobic turning. The resulting cell density, speed, and flux profiles are presented in the main text (Fig. 1h-j, solid gray curves). We show that the swimming speed of *C. reinhardtii* in bulk viscous media decays as a power law (Fig. 1f) with increasing viscosity, *η*:

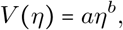

where a least squares fit gives *b* = −0.934 and *a* = 103, and *V* and *η* have units of *μ*m/s and cP, respectively. Combining this result with the known spatial variation of the viscosity, *η*(*x*), in our microfluidic gradient generating device – measured independently from microrheology experiments (Fig. 1g and Extended Data Fig. 1 and 5) – we obtain an expression for the swimming speed profile of the cells in a quasi-steady viscosity gradient:

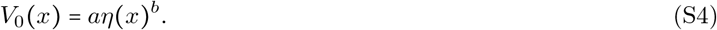

This analytical prediction (Fig. 1h, solid gray curves) agrees well with the local mean swimming speed profile of *C. reinhardtii* (Fig. 1h, solid colored curves), measured *in situ* for the viscotaxis experiments, and gives a high degree of confidence in the reliability of the predicted density profile below.

Next, we recognize that the product of the local density and speed (equation (S3)) should be constant throughout the channel under these conditions: *ρ*_0_(*x*)*V*_0_(*x*) = *k*. Combining equations (S4) and (S3) with this flux condition, we obtain the following expression for the predicted cell density profile:

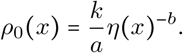

Because *ρ*_0_ is a probability density function, the normalization constant is defined by 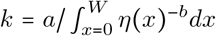, without any additional fitting parameters, and the expression for the predicted density profile (Fig. 1i, solid gray curves) is

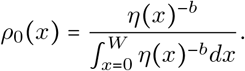

Following from the cell density, the predicted flux magnitude (i.e. a constant value; Fig. 1j, solid gray curves) in the viscosity gradient is

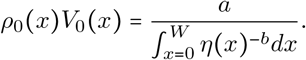

As described in the main text, *ρ*_0_ over-predicts the slope of the measured cell density profile, and the measured cell flux progressively deviates from the predicted constant value with increasing viscosity gradient. The strong agreement of the measured, *V*(*x*), and predicted, *V*_0_(*x*), velocity profiles combined with the deviations of the cell density and flux are indicative of a new phenomenon that depends on the magnitude of the viscosity gradient, namely the viscophobic reorientation described in the main text.

#### 4.2. Three-point force swimmer model

In the low Reynolds number regime, biflagellated swimming microalgae, such as *C. reinhardtii*, are commonly modeled using well-established three point force models [22, 23]. In particular, the thrust from the dual flagella are approximated by one point force each, while the drag from the body is approximated by a third point force or the Stokes drag on a spheroidal body. Here, we extend this idea for a biflagellated microswimmer in a viscosity gradient to corroborate our experimental observations (Fig. 2–3). The model considers a three point force model swimmer in a two-dimensional domain having an inhomogeneous viscosity landscape, *η*(**r**). In the analysis that follows, we approximate the drag and thrust forces as steady, which is equivalent to averaging over the flagellar beat cycle. This approximation is justified because the beat frequency of the flagella (~ 50 Hz) is at least two orders of magnitude faster than other relevant time scales, for example including the observed viscophobic turning rate, the effective rotational diffusion, and the travel time across the gradient. Furthermore, we assume the swimming dynamics do not affect the viscosity gradient (i.e. through mixing), which is consistent with other recent models [24–26]. In the analysis developed below, we illustrate that the two primary effects regulating the trajectory of a three-point force model swimmer in a viscosity gradient are (*i*) the orientation-dependent propulsive force imbalance between the two flagella and (*ii*) the coupling of rotational and translational motion of the cell body [24].

##### 4.2.1. Asymmetric flagellar propulsive force

For drag-based thrust in the Stokes flow regime, the effective propulsive force of each flagellum is proportional to the local viscosity, as well as the velocity and conformation of the flagellum. Thus, the force generated by a flagellum, *i*, is described as

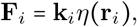

where **r**_*i*_ is the location of the center of thrust. The geometry and kinematics of the flagellum are encompassed in **k**_*i*_, which in the most general case is not necessarily aligned with the body axis. The center of thrust for a given flagellum can be expressed as

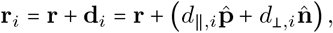

where **d**_*i*_ is the relative location from the body center (i.e. center of drag), **r**, and the cell body axis and its normal are 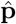 and 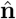, respectively. For linear viscosity gradients – or small separations of the center of thrust from the body center, compared to the viscosity gradient – the flagellar thrust is

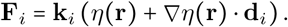

To focus on the effect of the viscosity gradient on the viscophobic turning behavior of the cells, we make the following simplifying assumptions in line with other recent simulations in viscosity gradients [26]: (*i*) The thrusting of the two flagella is symmetric and independent of the local viscosity, **k** = **k**_*i*_. (*ii*) The resultant thrusting is in the direction of the cell body, 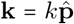. (*iii*) The centers of thrust of the two flagella are located lateral to the center of drag of the swimmer’s body, where 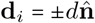 and *i* = 1, 2. The flagellar thrust forces become

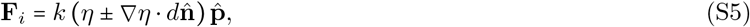

which also generate a torque about the center of the cell body

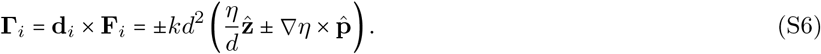

Thus, the forces and torques generated by the two flagella are not in general equal in the presence of a viscosity gradient.

##### 4.2.2. Rotational and translational coupling of swimmer body dynamics

The bodies of unicellular swimming microalgae – including *C. reinhardtii* used in the current work – are spherical to slightly prolate spheroids. Here, we model the dynamics of the swimmer body as a sphere of radius, *R*. Even for such highly symmetric bodies, a unique feature of their kinematics in a low Reynolds number viscosity gradient is the coupling of rotational and translational motion [24], which we quote below. The translational and rotational Stokes drag for a spherical body in a viscosity gradient are

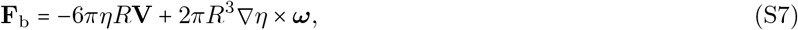

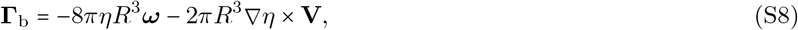

where **V** is the velocity (i.e. cell swimming speed), and ***ω*** is the angular rotation rate. Physically, if a sphere were to be dragged through a viscosity gradient at a velocity **V** in the direction perpendicular to the gradient, it would rotate at a speed ***ω***, but not if the force is applied in the viscosity gradient direction. This result is due to the differential viscous traction forces on either side of the body when moving perpendicular to the gradient, which ultimately cause it to rotate. In a similar fashion, if a sphere is rotated by an external torque, it will translate in the direction perpendicular to the gradient.

##### 4.2.3. Rotation rate of a swimmer in a viscosity gradient

Incorporating the flagellar thrust and cell body drag into a force and torque balance on the swimmer, we establish an expression for the rotation rate due to self-propulsion in a viscosity gradient, which we then compare to our experimental results in the main text. A torque balance on the three point force swimmer yields an expression in terms of the rotation rate:

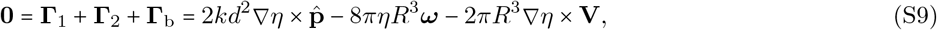

where the first and last terms on the right hand side represent the competition between viscophobic turning due to asymmetric flagellar thrust and viscophilic turning due to translational/rotational cell body coupling, respectively. Likewise, the force balance gives:

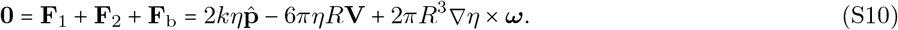

Recognizing that the rotation rate (equation (S9)) must be proportional to the viscosity gradient |***ω***| ∝ |▽*η*|, the final term of the force balance (equation (S10)) is second order in |▽*η*|. Thus, to first order in the viscosity gradient, swimmers largely move along their body axis with velocity

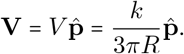

Inserting this result into equation (S9) and eliminating the flagellar thrust coefficient, *k*, the rotation rate of the swimmer, 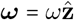, in a viscosity gradient becomes

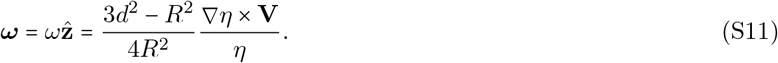

The prefactor depends solely on cell morphology and dictates whether swimmers are viscophobic or viscophilic. If the center of thrust of the flagella is significantly large compared to the cell radius, 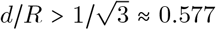, then cells will behave in a viscophobic manner and turn away from high viscosity zones. This condition is consistent with the typical geometry of *C. reinhardtii* and our experimental observations of cell turning (Fig. 2–3). Conservatively, for a typical cell radius [12], R ≈ 5 *μ*m, flagellar length, *ℓ* ≈ 11 *μ*m, and flagellar center of thrust, *d* ≈ *ℓ*/2, wild-type *C. reinhardtii* are generally expected to exhibit viscophobic behavior, *d*/*R* ≈ 1.1 > 0.577. Furthermore, the prefactor in equation (S11) reduces to ≈ 0.66, which depends strongly on flagellar length and is similar in magnitude to the factor 1/2 in recent squirmer models [26].

For a one-dimensional viscosity profile, *η*(*x*), and a viscosity dependent swimming velocity, 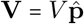, the viscophobic rotation rate is

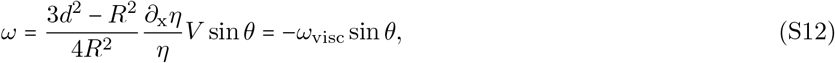

where *ω*_visc_ is the viscophobic turning rate amplitude, *θ* is the swimming direction, 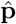. The geometrical prefactor above depends upon the particular microhydrodynamics of the cell motility, which is beyond the scope of this work. However, this analysis allows us to make quantitative predictions on the scaling of the swimmer turning rate amplitude, which is linearly proportional to the swimming speed, *ω*_visc_ ∝ *V*, as might be expected for low Reynolds number locomotion. Importantly, we also find that the turning rate scales with *ω*_visc_ ∝ |▽*η*|/*η*, which agrees with our experiments (Fig. 2) and is also consistent with recent predictions for squirmer models [26].

#### 4.3. Viscophobic turning trajectories

Experimentally measured trajectories of swimming *C. reinhardtii* were demonstrated to turn away from regions of high viscosity and toward regions of low viscosity in the presence of a viscosity gradient (Fig. 2a,b). In particular, cells starting with initial orientation *θ*_0_ = ±*π*/2 in the center of the maximum viscosity gradient (▽*η* = 7.2 × 10^−3^ cP·*μ*m^−1^), reveal smoothly curved trajectories stemming from viscophobic turning (Fig. 2b). Here, we examine the equations of motion for viscophobic turning and develop an analytical expression for the expected cell trajectories (Fig. 2b, black curve), which show strong agreement with experiments. The equations of motion for a cell with local swimming speed, *V*, become

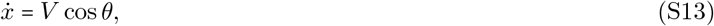

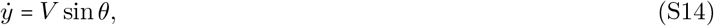

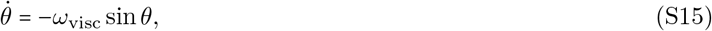

where *θ* is the cell orientation with *θ* = 0 being in the direction of decreasing viscosity, and *ω*_visc_ is the viscophobic turning rate amplitude (Fig. 2c,d). Substituting *u* = tan(*θ*/2), we write

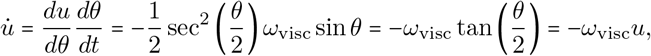

where the over dot indicates a time derivative. Integrating the above expression yields 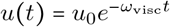 or 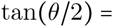 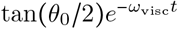. This allows us to rewrite the equations of motion as

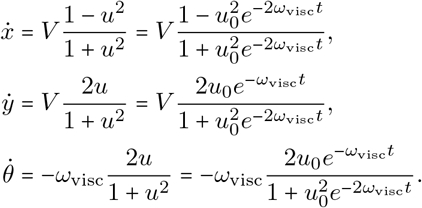

Integrating 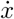 and 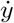 gives a parametric expression for the cell trajectory in *x*(*t*) and *y*(*t*) as

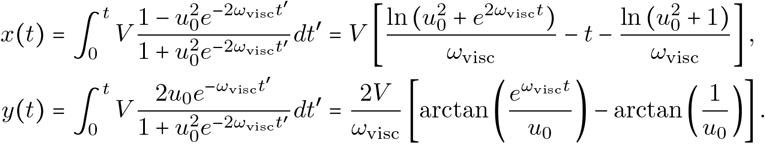

Finally, if we consider initial orientations of the cells in the direction perpendicular to the gradient, *θ*_0_ = ±*π*/2 and *u*_0_ = tan(*θ*_0_/2) = ±1, the trajectory becomes:

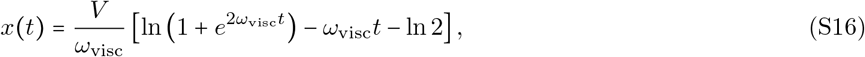

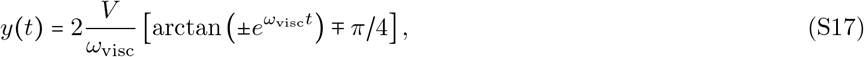

which is shown in Fig. 2b (solid black curve). While this relatively simple analysis omits the stochasticity of cell motility and spatial variations in swimming speed, the model largely captures the shape of the observed cell trajectories in the viscosity gradient.

### 5. STOCHASTIC DYNAMICS OF VISCOPHOBIC TURNING MOTILITY

#### 5.1. Langevin simulation

##### 5.1.1. Langevin equations of motion

Microswimmers such as *C. reinhardtii* are subject to stochasticity in their motility stemming from both thermal motion and noise in their flagellar propulsion. In particular, noise in the cell orientation dominates the effects on self-transport compared to translation [27, 28]. We model these effects as a persistent random walk by the addition of a noise term in the orientation dynamics and obtain the Langevin equations of motion

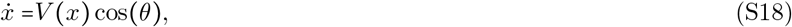

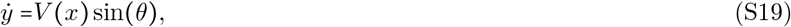

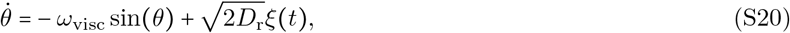

where *D*_r_ is the effective rotational diffusion coefficient of the cells, and *ξ*(t) is a Gaussian noise term. The rotational diffusion coefficient is taken to be a constant, as our experiments indicate that it does not vary significantly over the range of viscosities explored in this work (Extended Data Fig. 7 and Methods). Here, we allow the swimming speed, *V*(*x*), to vary spatially, which captures the main effect of viscous slow-down on cell translational motion. For simplicity, we take the cell speed profile to be linear across the channel *V*(*x*) = (*V*_max_ − Δ*V*) + Δ*V*_*x*_/*W*, where *V*_max_ and *V*_min_ are the cell speed at *x* = *W* and *x* = 0, respectively, and Δ*V* = *V*_max_ − V_min_. We also define the dimensionless variables

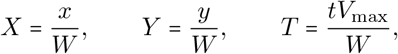

which lead to the following dimensionless coefficients

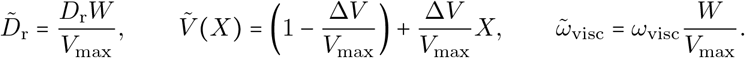

Finally, substituting into the Langevin equations of motion, we obtain

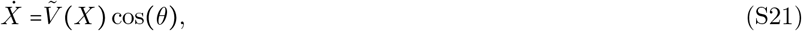

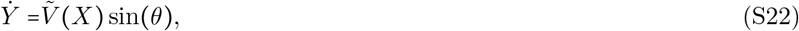

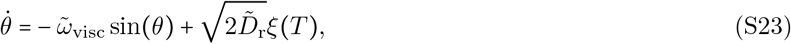

where the over dot now indicates derivatives with respect to dimensionless time, T. Thus, the Langevin equation dynamics are ultimately dictated by the three dimensionless parameters 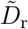, Δ*V*/*V*_max_, and 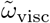. Due to the weak variation of rotational diffusivity, 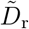 was held constant. Simulations were performed as described in the Methods and as a function of Δ*V*/*V*_max_ and 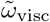, where the results are shown in Extended Data Fig. 2–3 and in the main text (Fig. 3).

##### 5.1.2. Measured cell-surface interaction and Langevin simulation boundary condition

Swimming cells are well-known to accumulate near solid surfaces due to the nature of their steric and hydrodynamic interactions with boundaries [29, 30]. By parameterizing boundary interactions as scattering events, previous experimental work has demonstrated that random surface scattering by *C. reinhardtii* is biased toward small angles [30], *θ*_out_, relative to the surface parallel (Fig. S4a, inset). To definitively establish the relevant boundary condition for our Langevin simulation, we directly measure the scattering angle distribution of wild-type *C. reinhardtii* (Fig. S4a) from our control experiments in the absence of a viscosity gradient (▽*η* = 0; see also Fig. 1). The measured scattering angle distribution is in excellent agreement with previous work, and it is fitted by a non-parametric kernel estimator (MATLAB; Fig. S4a, red curve), providing a smooth PDF to efficiently sample in our simulations.

To implement the empirically measured boundary condition in Langevin simulations, we treat the scattering events as rapid reorientations of the simulated cells toward randomly selected angles, *θ*_out_, chosen from the measured scattering distribution. Upon contacting the boundary (*X* ≤ 0 or *X* ≥ 1), cells are rotated with an additional angular velocity, Ω_wall_:

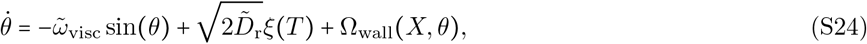

where Ω_wall_ aligns the cells toward a randomly selected *θ*_out_ oriented inside the channel (Fig. S4a, red curve). This reorientation is set by a paramagnetic-like potential [31], which is defined as

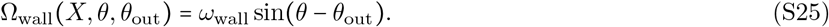

The constant *ω*_wall_ is set to 1000 s^−1^ to ensure that cells reorient to their randomly selected scattering angle within ≈ 0.005 s. The density profiles resulting from Langevin simulations for the control (Fig. 1i, dashed black) exhibit excellent agreement with the experiments (Fig. 1i, solid black) and accurately capture the mild surface accumulation. Furthermore, we expect no coupling between the viscophobic turning phenomenon discussed in the main text and the observed cell-surface interactions, which are confined to the very near-wall region. The simulations confirm this notion and accurately capture the measured cell densities across all tested viscosity gradients (Fig. 1i, solid and dashed colored lines)

#### 5.2. Semi-empirical relation for turning rate and swimming speed contrast

Langevin simulations facilitate the arbitrary exploration of the parameter space of *ω*_visc_ and Δ*V*/*V*_max_ (Fig. 3c). However, experimentally, both parameters directly depend upon the viscosity gradient (Figs. 1f and 2d), which is formed by two fluids of high and low viscosity, initially injected into the test channel. During the experiment time, the extrema of the channel evolve to corresponding viscosities *η*_max_ and *η*_min_ (Fig. 1g), whose ratio parametrically controls the trajectory of experiments in the parameters space (Fig. 3c). Here, we develop a semi-empirical relationship for the evolution of *ω*_visc_ with Δ*V*/*V*_max_ based on this control parameter *η*_max_/*η*_min_. The swimming speed contrast is evaluated from the power-law relationship (Fig. 1f) of the swimming speed with viscosity, *V* = a*η*^−b^:

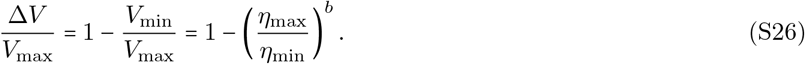

Similarly, the observed linear relationship for viscophobic rotation rate (Fig. 2d; equation S12) can be written in terms of the approximate viscosity gradient ▽*η* ≈ (*η*_max_ − *η*_min_)/*W* and mean viscosity *η*_0_ ≈ *β*(*η*_max_ + *η*_min_)/2:

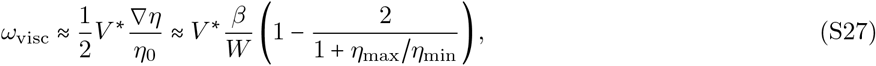

where *V*\* = 66 *μ*m/s is based on the fit in Fig. 2d and *β* = 1.7 ± 0.1 accounts for the observed magnitude of the gradient in the central region of the channel relative to the slightly sigmoidal shape in our experiments (Fig. 1g). Thus, equations (S26) and (S27) form a parametric set of equations in *η*_max_/*η*_min_, which well represent the experimental observations and matching Langevin simulations (Fig. 3c, black curve).

#### 5.3. Fokker-Planck model of viscophobic reorientation dynamics

To complement the experiments and Langevin simulations discussed above, we also employ a simplified one-dimensional Fokker-Planck (FP) model. The FP model takes the form of a rotational advection diffusion equation to describe the joint probability density of orientation and time, *p*(*θ*,*t*). The swimming cells are subjected to an orientation dependent rotation rate due to viscophobic turning motility, and they experience random orientational fluctuations parameterized by rotational diffusion:

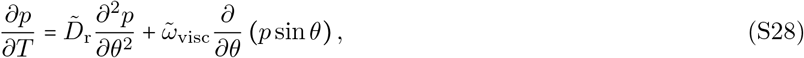

where 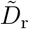 and 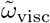 are constants, as above. The equation is solved using a second-order finite difference scheme for various 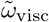, corresponding to experimental conditions that include both control experiments (▽*η* = 0 cP·*μ*m^−1^; Extended Data Fig. 2c) and maximum viscosity gradient conditions (▽*η* = 7.2 × 10^−3^ cP·*μ*m^−1^; Extended Data Fig. 2f). As with the Langevin simulations, the turning rate amplitude and rotational diffusion parameters are taken directly from experimental measurements, and thus, there are no fitting parameters. The result is viewed as a conditional probability density, p(*θ*|*t*), that illustrates the time evolution of the cell orientation distribution (Extended Data Fig. 2). Physically, the cell rotation rate in the viscosity gradient drives a condensation, or narrowing, of the orientation distribution toward the low viscosity direction (*θ* = 0). This effect is in competition with random cell reorientation, which tends to widen the distribution. The results of the FP theory were compared to both experiments (Extended Data Fig. 2a,d) and Langevin simulations (Extended Data Fig. 2b,e), which remain accurate (Fig. 3b) when the cells are isolated far from the channel walls.

### 6. SUPPLEMENTARY FIGURES

**Fig. S1.**
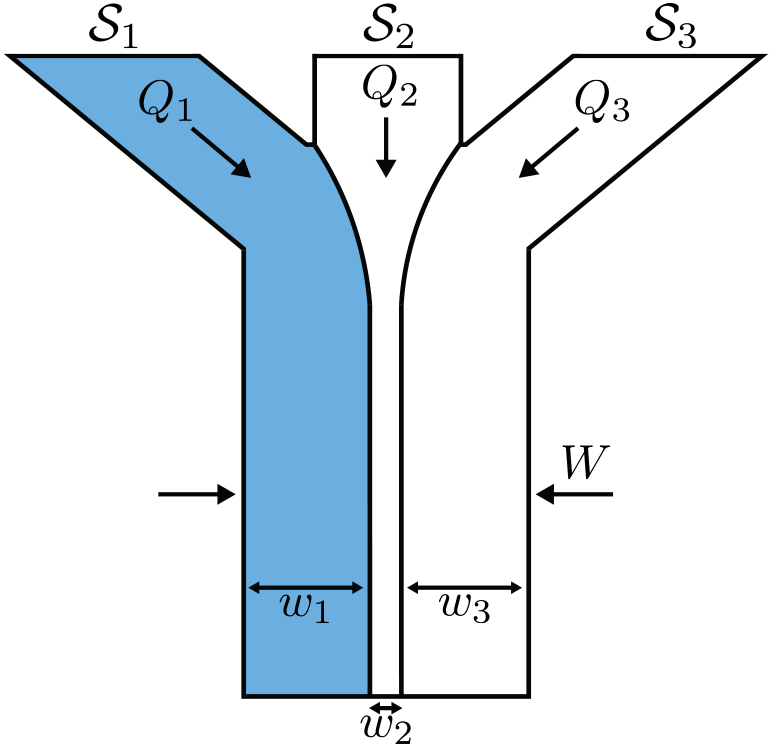
Microfluidic viscosity gradient generation. Schematic of the junction and test section of a microfluidic viscosity gradient generation device, where inlets containing a viscous solution of PEO polymer in M1 media (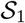; left), *C. reinhardtii* cell suspension in M1 (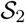; center), and M1 media (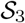; right) meet. The flow rates *Q*_1_, *Q*_2_, and *Q*_3_ are adjusted for solutions having viscosities *η*_1_, *η*_2_, and *η*_3_ to maintain the initial stratification of these streams in the main channel of *w*_1_ = *w*_3_ = 4*W*/9 and *w*_2_ = *W*/9. Background is qualitative and illustrates initial stratification of viscous fluid (blue), cell suspension, and media.

**Fig. S2.**
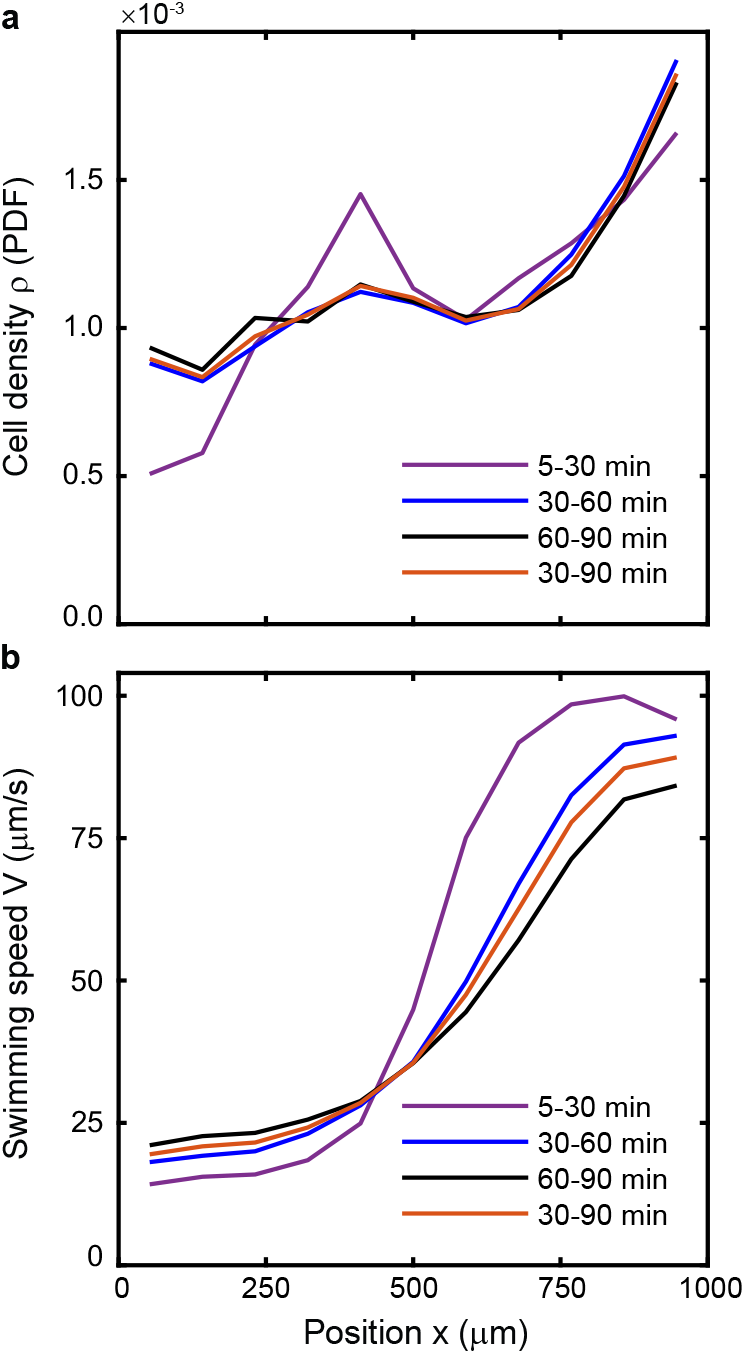
Comparison of measured local cell density and swimming speed for various assay time intervals. A comparison of wild-type *C. reinhardtii* at different time intervals throughout the viscotaxis experiment in the maximum viscosity gradient tested ▽*η* = 7.2 × 10^−3^ cP·*μ*m^−1^). **a**, The distribution of *C. reinhardtii* cell density between 30-60 min and 60-90 min intervals remains relatively constant, with a mean percent difference of 1.84% and 2.33%, respectively, when compared to the full analysis time interval of 30-90 min. The time interval 5-30 min, during which time the viscosity profile is strongly evolving (Extended Data Fig. 1), is shown for reference, **b**, The distribution of the cell’s local mean swimming speed exhibits similarly small deviations for the 30-60 min and 60-90 min time intervals, having a mean percent error of 5.08% and 6.08%, respectively, when compared to the full analysis time interval of 30-90 min.

**Fig. S3.**
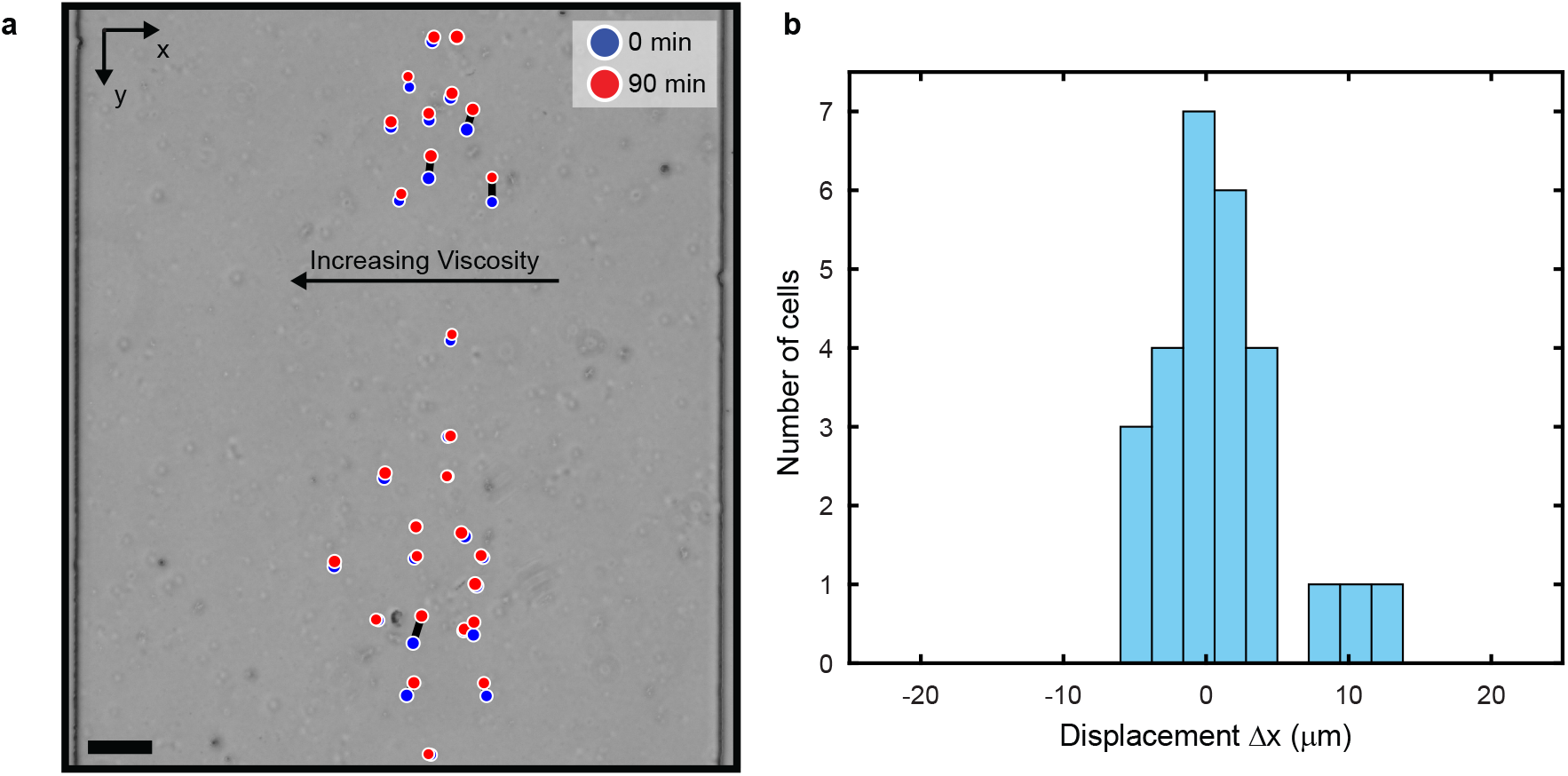
Control experiments exhibit no discernible diffusiophoretic drift of non-motile wild-type *C. reinhardtii.* **a**, Viscotaxis control experiments using (fixed) non-motile cells in the maximum viscosity gradient tested (▽*η* = 7.2 × 10^−3^ cP·*μ*m^−1^) show no significant observable displacement over the course of a 90 min experiment under identical conditions to Fig. 1h,i (Scale bar, 100 *μ*m). **b**, Maximum cell displacements in the viscosity gradient direction from **a** are minute (on the order of ≈ 1 cell diameter or 1% of the channel width) and consistent with Brownian diffusion of the non-motile cells.

**Fig. S4.**
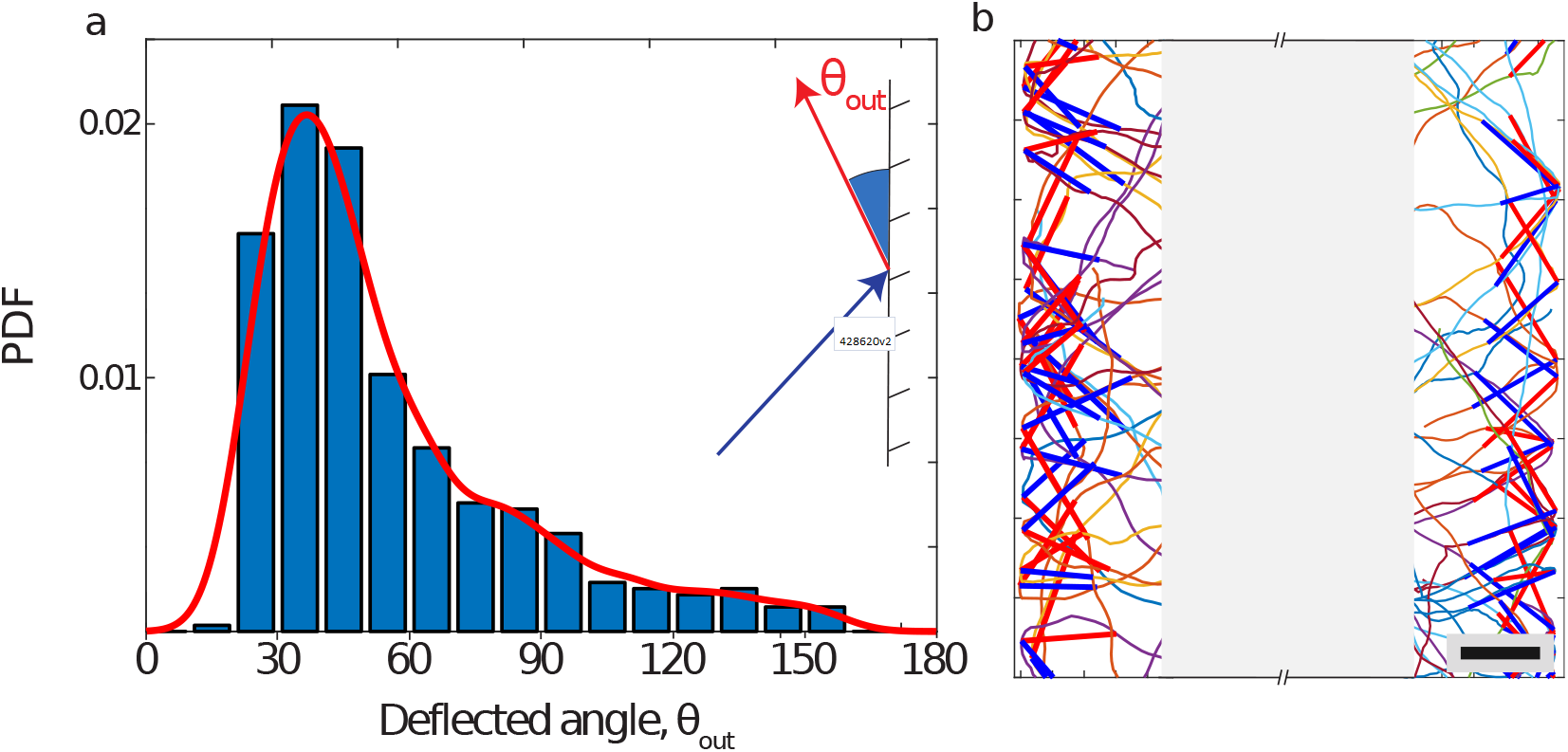
Cell scattering at solid boundaries. **a**, Measured outgoing scattering angle distribution (blue) for wild-type *C. reinhardtii* encountering the microfluidic channel walls (396 events) in the absent of a viscosity gradient (control, ▽*η* = 0). Red curve is the fitted non-parametric kernel distribution (MATLAB) used to randomly generate scattering angles in Langevin simulations. Inset: Definition of scattering angle, *θ*_out_, relative to the wall-parallel direction. b, Sample measured cell scattering events from cell tracks (multicolored curves). Blue and red segments show detected incoming and outgoing cells. Channel center is masked in light gray. Scale bar, 50 *μ*m.

## References

1 Wild, C. et al. Coral mucus functions as an energy carrier and particle trap in the reef ecosystem. Nature 428, 66–70 (2004).

2 Wheeler, K. M. et al. Mucin glycans attenuate the virulence of *Pseudomonas aeruginosa* in infection. Nat. Microbiol. 4, 2146–2154 (2019).

3 Suarez, S. S. & Pacey, A. A. Sperm transport in the female reproductive tract. Hum. Reprod. Update 12, 23–37 (2006).

4 Hall-Stoodley, L., Costerton, J. W. & Stoodley, P. Bacterial biofilms: from the natural environment to infectious diseases. Nat. Rev. Microbiol. 2, 95–108 (2004).

5 Swidsinski, A. et al. Viscosity Gradient Within the Mucus Layer Determines the Mucosal Barrier Function and the Spatial Organization of the Intestinal Microbiota. Inflamm. Bowel Dis. 13, 963–970 (2007).

6 Petrino, M. G. & Doetsch, R. N. ‘Viscotaxis’, a New Behavioural Response of *Leptospira interrogans* (*biflexa*) Strain b16. J. Gen. Microbiol. 109, 113–117 (1978).

7 Daniels, M. J., Longland, J. M. & Gilbart, J. Aspects of Motility and Chemotaxis in Spiroplasmas. J. Gen. Microbiol. 118, 429–436 (1980).

8 Sherman, M. Y., Timkina, E. & Glagolev, A. Viscosity taxis in *Escherichia coli*. FEMS Microbiol. Lett. 13, 137–140 (1982).

9 Schnitzer, M. J. Theory of continuum random walks and application to chemotaxis. Phys. Rev. E 48, 2553–2568 (1993).

10 Frangipane, G. et al. Dynamic density shaping of photokinetic *E. coli*. Elife 7, e36608 (2018).

11 Liebchen, B., Monderkamp, P., Ten Hagen, B. & Löwen, H. Viscotaxis: Microswimmer Navigation in Viscosity Gradients. Phys. Rev. Lett. 120, 208002 (2018).

12 Datt, C. & Elfring, G. J. Active Particles in Viscosity Gradients. Phys. Rev. Lett. 123, 158006 (2019).

13 Qin, B., Gopinath, A., Yang, J., Gollub, J. P. & Arratia, P. E. Flagellar Kinematics and Swimming of Algal Cells in Viscoelastic Fluids. Sci. Rep. 5, 9190 (2015).

14 Stocker, R. Marine Microbes See a Sea of Gradients. Science 338, 628–633 (2012).

15 Cavicchioli, R. et al. Scientists’ warning to humanity: microorganisms and climate change. Nat. Rev. Microbiol. 17, 569–586 (2019).

16 Berg, H. C. & Brown, D. A. Chemotaxis in *Escherichia coli* analysed by Three-dimensional Tracking. Nature 239, 500–504 (1972).

17 Jékely, G. et al. Mechanism of phototaxis in marine zooplankton. Nature 456, 395–399 (2008).

18 Durham, W. M., Kessler, J. O. & Stocker, R. Disruption of Vertical Motility by Shear Triggers Formation of Thin Phytoplankton Layers. Science 323, 1067–1070 (2009).

19 Sengupta, A., Carrara, F. & Stocker, R. Phytoplankton can actively diversify their migration strategy in response to turbulent cues. Nature 543, 555–558 (2017).

20 Blakemore, R. P. Magnetotactic bacteria. Science 190, 377–379 (1975).

21 Guadayol, Ò. et al. Microrheology reveals microscale viscosity gradients in planktonic systems. Proc. Natl Acad. Sci. USA 118, e2011389118 (2021).

22 Lai, S. K., Wang, Y. Y., Wirtz, D. & Hanes, J. Micro- and macrorheology of mucus. Adv. Drug Deliv. Rev. 61, 86–100 (2009).

23 Jatkar, A. A. et al. Measuring mucus thickness in reef corals using a technique devised for vertebrate applications. Mar. Biol. 157, 261–267 (2010).

24 Atuma, C., Strugala, V., Allen, A. & Holm, L. The adherent gastrointestinal mucus gel layer: thickness and physical state in vivo. Am. J. Physiol. Liver Physiol. 280, G922–G929 (2001).

25 Stabili, L., Schirosi, R., Parisi, M. G., Piraino, S. & Cammarata, M. The Mucus of *Actinia equina* (Anthozoa, Cnidaria): An Unexplored Resource for Potential Applicative Purposes. Mar. Drugs 13, 5276–5296 (2015).

26 Cone, R. A. Barrier properties of mucus. Adv. Drug Deliv. Rev. 61, 75–85 (2009).

27 Martinez, V. A. et al. Flagellated bacterial motility in polymer solutions. Proc. Natl Acad. Sci. USA 111, 17771–17776 (2014).

28 Shoele, K. & Eastham, P. S. Effects of nonuniform viscosity on ciliary locomotion. Phys. Rev. Fluids 3, 043101 (2018).

29 Harris, E. H. The Chlamydomonas Sourcebook: Introduction to Chlamydomonas and its laboratory use, vol. 1 (Academic Press, 2009).

30 Goldstein, R. E. Green Algae as Model Organisms for Biological Fluid Dynamics. Annu. Rev. Fluid Mech. 47, 343–375 (2015).

31 Kantsler, V., Dunkel, J., Polin, M. & Goldstein, R. E. Ciliary contact interactions dominate surface scattering of swimming eukaryotes. Proc. Natl Acad. Sci. USA 110, 1187–1192 (2013).

32 Cates, M. E. Diffusive transport without detailed balance in motile bacteria: does microbiology need statistical physics? Reports Prog. Phys. 75, 042601 (2012).

33 Polin, M., Tuval, I., Drescher, K., Gollub, J. P. & Goldstein, R. E. *Chlamydomonas* Swims with Two “Gears” in a Eukaryotic Version of Run-and-Tumble Locomotion. Science 325, 487–490 (2009).

34 Geyer, V. F., Jölicher, F., Howard, J. & Friedrich, B. M. Cell-body rocking is a dominant mechanism for flagellar synchronization in a swimming alga. Proc. Natl Acad. Sci. USA 110, 18058–18063 (2013).

35 Lauga, E. & Powers, T. R. The hydrodynamics of swimming microorganisms. Reports Prog. Phys. 72, 096601 (2009).

36 Kuchka, M. R. & Jarvik, J. W. Short-Flagella Mutants of *Chlamydomonas reinhardtii*. Genetics 115, 685–691 (1987).

37 Oppenheimer, N., Navardi, S. & Stone, H. A. Motion of a hot particle in viscous fluids. Phys. Rev. Fluids 1, 014001 (2016).

38 Waisbord, N. & Guasto, J. S. Peculiar polygonal paths. Nat. Phys. 14, 1161–1162 (2018).

39 Rupprecht, J. F., Waisbord, N., Ybert, C., Cottin-Bizonne, C. & Bocquet, L. Velocity Condensation for Magnetotactic Bacteria. Phys. Rev. Lett. 116, 168101 (2016).

40 Sager, R. & Granick, S. Nutritional studies with *Chlamydomonas reinhardi*. Ann. N. Y. Acad. Sci. 56, 831–838 (1953).

41 Ebagninin, K. W., Benchabane, A. & Bekkour, K. Rheological characterization of poly(ethylene oxide) solutions of different molecular weights. J. Colloid Interface Sci. 336, 360–367 (2009).

42 Grigorescu, G. & Kulicke, W.-M. Prediction of Viscoelastic Properties and Shear Stability of Polymers in Solution, 1–40 (Springer, Berlin, Heidelberg, 2000).

43 Xia, Y. & Whitesides, G. M. SOFT LITHOGRAPHY. Annu. Rev. Mater. Sci. 28, 153–184 (1998).

44 Devanand, K. & Selser, J. C. Polyethylene oxide does not necessarily aggregate in water. Nature 343, 739–741 (1990).

45 Waigh, T. A. Microrheology of complex fluids. Reports Prog. Phys. 68, 685–742 (2005).

46 Kreis, C. T., Le Blay, M., Linne, C., Makowski, M. M. & Bäumchen, O. Adhesion of *Chlamydomonas* microalgae to surfaces is switchable by light. Nat. Phys. 14, 45–49 (2018).

47 Rafaï, S., Jibuti, L. & Peyla, P. Effective Viscosity of Microswimmer Suspensions. Phys. Rev. Lett. 104, 098102 (2010).

48 Cheezum, M. K., Walker, W. F. & Guilford, W. H. Quantitative Comparison of Algorithms for Tracking Single Fluorescent Particles. Biophys. J. 81, 2378–2388 (2001).

49 Crocker, J. C. & Grier, D. G. Methods of Digital Video Microscopy for Colloidal Studies. J. Colloid Interface Sci. 179, 298–310 (1996).

50 Ouellette, N. T., Xu, H. & Bodenschatz, E. A quantitative study of three-dimensional Lagrangian particle tracking algorithms. Exp. Fluids 40, 301–313 (2006).

51 Crenshaw, H. C. A New Look at Locomotion in Microorganisms: Rotating and Translating. Am. Zool. 36, 608–618 (1996).

52 Howse, J. R. et al. Self-Motile Colloidal Particles: From Directed Propulsion to Random Walk. Phys. Rev. Lett. 99, 48102 (2007).

53 Rusconi, R., Guasto, J. S. & Stocker, R. Bacterial transport suppressed by fluid shear. Nat. Phys. 10, 212–217 (2014).

54 Dehkharghani, A., Waisbord, N., Dunkel, J. & Guasto, J. S. Bacterial scattering in microfluidic crystal flows reveals giant active taylor–aris dispersion. Proc. Natl Acad. Sci. USA 116, 11119–11124 (2019).

55 Korson, L., Drost-Hansen, W. & Millero, F. J. Viscosity of water at various temperatures. J. Phys. Chem. 73, 34–39 (1969).

## References

[1] Sager, R. & Granick, S. Nutritional studies with *Chlamydomonas reinhardi*. Ann. N. Y. Acad. Sci. 56, 831–838 (1953).

[2] Devanand, K. & Selser, J. C. Polyethylene oxide does not necessarily aggregate in water. Nature 343, 739–741 (1990).

[3] Martinez, V. A. et al. Flagellated bacterial motility in polymer solutions. Proc. Natl Acad. Sci. USA 111, 17771–17776 (2014).

[4] Kirby, B. J. Micro-and nanoscale fluid mechanics: transport in microfluidic devices (Cambridge University Press, 2010).

[5] Rafaï, S., Jibuti, L. & Peyla, P. Effective Viscosity of Microswimmer Suspensions. Phys. Rev. Lett. 104, 098102 (2010).

[6] Gardel, M. L., Valentine, M. T. & Weitz, D. A. Microrheology. In Breuer, K. S. (ed.) Microscale Diagnostic Tech., 1–49 (Springer, Berlin, Heidelberg, 2005).

[7] Liu, J. et al. Microrheology Probes Length Scale Dependent Rheology. Phys. Rev. Lett. 96, 118104 (2006).

[8] Waigh, T. A. Microrheology of complex fluids. Reports Prog. Phys. 68, 685–742 (2005).

[9] Ebagninin, K. W., Benchabane, A. & Bekkour, K. Rheological characterization of poly(ethylene oxide) solutions of different molecular weights. J. Colloid Interface Sci. 336, 360–367 (2009).

[10] Goldstein, R. E. Green Algae as Model Organisms for Biological Fluid Dynamics. Annu. Rev. Fluid Mech. 47, 343–375 (2015).

[11] Harris, E. H. *Chlamydomonas* as a model organism. Annu. Rev. Plant Physiol. Plant Mol. Biol. 52, 363–406 (2001).

[12] Harris, E. H. The Chlamydomonas Sourcebook: Introduction to Chlamydomonas and its laboratory use, vol. 1 (Academic Press, 2009).

[13] Kuchka, M. R. & Jarvik, J. W. Short-Flagella Mutants of *Chlamydomonas reinhardtii*. Genetics 115, 685–691 (1987).

[14] Qin, B., Gopinath, A., Yang, J., Gollub, J. P. & Arratia, P. E. Flagellar Kinematics and Swimming of Algal Cells in Viscoelastic Fluids. Sci. Rep. 5, 9190 (2015).

[15] Berg, H. C. & Turner, L. Movement of microorganisms in viscous environments. Nature 278, 349–351 (1979).

[16] Schneider, W. R. & Doetsch, R. N. Effect of Viscosity on Bacterial Motility. J. Bacteriol. 117, 696–701 (1974).

[17] Berg, H. C. & Brown, D. A. Chemotaxis in *Escherichia coli* analysed by Three-dimensional Tracking. Nature 239, 500–504 (1972).

[18] Garren, M. et al. A bacterial pathogen uses dimethylsulfoniopropionate as a cue to target heat-stressed corals. ISME J. 8, 999–1007 (2014).

[19] Moran, J. L. & Posner, J. D. Phoretic Self-Propulsion. Annu. Rev. Fluid Mech. 49, 511–540 (2017).

[20] Leptos, K. C., Guasto, J. S., Gollub, J. P., Pesci, A. I. & Goldstein, R. E. Dynamics of Enhanced Tracer Diffusion in Suspensions of Swimming Eukaryotic Microorganisms. Phys. Rev. Lett. 103, 198103 (2009).

[21] Schnitzer, M. J. Theory of continuum random walks and application to chemotaxis. Phys. Rev. E 48, 2553–2568 (1993).

[22] Drescher, K., Goldstein, R. E., Michel, N., Polin, M. & Tuval, I. Direct Measurement of the Flow Field around Swimming Microorganisms. Phys. Rev. Lett. 105, 168101 (2010).

[23] Bennett, R. R. & Golestanian, R. A steering mechanism for phototaxis in *Chlamydomonas*. J. R. Soc. Interface 12, 20141164 (2015).

[24] Oppenheimer, N., Navardi, S. & Stone, H. A. Motion of a hot particle in viscous fluids. Phys. Rev. Fluids 1, 014001 (2016).

[25] Liebchen, B., Monderkamp, P., Ten Hagen, B. & Löwen, H. Viscotaxis: Microswimmer Navigation in Viscosity Gradients. Phys. Rev. Lett. 120, 208002 (2018).

[26] Datt, C. & Elfring, G. J. Active Particles in Viscosity Gradients. Phys. Rev. Lett. 123, 158006 (2019).

[27] Howse, J. R. et al. Self-Motile Colloidal Particles: From Directed Propulsion to Random Walk. Phys. Rev. Lett. 99, 48102 (2007).

[28] Rusconi, R., Guasto, J. S. & Stocker, R. Bacterial transport suppressed by fluid shear. Nat. Phys. 10, 212–217 (2014).

[29] Drescher, K., Dunkel, J., Cisneros, L. H., Ganguly, S. & Goldstein, R. E. Fluid dynamics and noise in bacterial cell–cell and cell–surface scattering. Proc. Natl Acad. Sci. USA 108, 10940–10945 (2011).

[30] Kantsler, V., Dunkel, J., Polin, M. & Goldstein, R. E. Ciliary contact interactions dominate surface scattering of swimming eukaryotes. Proc. Natl Acad. Sci. USA 110, 1187–1192 (2013).

[31] Dehkharghani, A., Waisbord, N., Dunkel, J. & Guasto, J. S. Bacterial scattering in microfluidic crystal flows reveals giant active taylor–aris dispersion. Proc. Natl Acad. Sci. USA 116, 11119–11124 (2019).

